# High-throughput optimized prime editing mediated endogenous protein tagging for pooled imaging of protein localization

**DOI:** 10.1101/2024.09.16.613361

**Authors:** Henry M. Sanchez, Tomer Lapidot, Ophir Shalem

**Affiliations:** Center for Cellular and Molecular Therapeutics, Children’s Hospital of Philadelphia, Philadelphia, PA 19104, USA; Department of Genetics, Perelman School of Medicine, University of Pennsylvania, Philadelphia, PA 19104, USA; Department of Bioengineering, University of Pennsylvania, Philadelphia, PA 19104, USA

## Abstract

The subcellular organization of proteins carries important information on cellular state and gene function, yet currently there are no technologies that enable accurate measurement of subcellular protein localizations at scale. Here we develop an approach for pooled endogenous protein tagging using prime editing, which coupled with an optical readout and sequencing, provides a snapshot of proteome organization in a manner akin to perturbation-based CRISPR screens. We constructed a pooled library of 17,280 pegRNAs designed to exhaustively tag 60 endogenous proteins spanning diverse localization patterns and explore a large space of genomic and pegRNA design parameters. Pooled measurements of tagging efficiency uncovered both genomic and pegRNA features associated with increased efficiency, including epigenetic states and interactions with transcription. We integrate pegRNA features into a computational model with predictive value for tagging efficiency to constrain the design space of pegRNAs for large-scale peptide knock-in. Lastly, we show that combining in-situ pegRNA sequencing with high-throughput deep learning image analysis, enables exploration of subcellular protein localization patterns for many proteins in parallel following a single pooled lentiviral transduction, setting the stage for scalable studies of proteome dynamics across cell types and environmental perturbations.

## Introduction

System-level understanding of cellular state and gene function requires technologies for visualizing proteins in their endogenous regulatory context at scale. Protein tagging provides a specific and generic handle to target and visualize proteins in living cells. Arrayed generation of endogenously tagged cell lines has demonstrated that large imaging datasets of protein localization patterns, coupled with deep learning image analysis, provides unique and crucial insights into cellular organization and protein function^1^. Since these approaches still rely on the delivery of gene-specific reagents (e.g., homology-dependent repair templates) in an arrayed format, they remain labor intensive and limited in scale. Adapting such technologies to a pooled format, such that imaging of hundreds to thousands of proteins could be done in a manner similar to a CRISPR screen, will enable unprecedented insight into changes in proteome organization in response to genetic and environmental perturbations.

As endogenous tagging through homology-dependent repair (HDR), requires the delivery of both a targeted nuclease and donor insertion template, lack of tools for paired delivery of these two components has limited the development of methods for pooled endogenous protein tagging. One way to avoid the need for gene-specific donors is to use a generic donor and rely on non-homologous end joining (NHEJ) as the DNA repair pathway. Such approach has been used for pooled tagging, yet for terminus tagging it is only compatible with the C terminus with a limited choice of sgRNAs, resulting in low success rates and likely incorporation of coding mutations^2,3^. To combine generic donors with large flexibility in tag site and choice of sgRNAs, we have previously developed scalable intron tagging by introducing a synthetic exon within introns^4^. Indeed, this approach has been used successfully to isolate a large array of endogenously tagged cell clones from a tagged pool^5^. We have recently expanded this approach to study unfolded proteins in an unbiased manner^6^ and search for proteins with degradation function through induced proximity^7^, demonstrating the utility of pooled protein tagging for biological discovery. Nonetheless, as these NHEJ mediated approaches: require separate delivery of DNA donor templates at high concentration, insert tags in random orientation, require multiple libraries to tag three possible coding frames and carry chromosomal risks associated with double-strand breaks (DSBs) – there remains a technology gap to advance pooled tagging as a standard CRISPR screening modality.

The development of prime editing has introduced a versatile method for genome editing that facilitates insertion mutagenesis using a single gene-specific reagent by encoding the insertion template within an extension of the sgRNA^8,9^. Prime editing works by using a programmable DNA nickase such as SpCas9(H840A) fused to a reverse transcriptase such as MMLV-RT. This Cas9-RT fusion is called a prime editor and is guided to a target site of interest by a prime editing guide RNA (pegRNA) encoding the desired edit in its 3’ extension. When the prime editor-pegRNA complex binds to the target DNA it nicks the non-target strand allowing the nicked DNA to function as a primer that binds to the primer-binding site (PBS) at the end of the 3’ extension. This hybridization event is followed by reverse transcription of the template in the rest of the extension. The edited DNA replaces the original DNA following resolution of the heteroduplex containing edited and unedited strands. While still limited in insertion size, such approach has many advantages for pooled tagging as it enables the precise insertion of peptides in both N and C termini without generating DSBs, uses an RNA template that can be delivered in lentiviral vectors and amplified through transcription, has minimal off-target editing and works across diverse cell types.

Prime editing has been used successfully for pooled genome editing aimed at modeling prime editing efficiency^10–13^ and high-throughput investigation of variant effects^14,15^, yet its application to pooled insertions at multiple unique loci across the genome has been limited – likely due to generally low insertion efficiencies associated with single flap prime editing as well as lack of feature-complete pegRNA design models for larger insertions with sought-after functions. Here we focused on developing a scalable and modular approach for pooled N and C terminus tagging using prime editing and a split fluorescent protein system, while laying the groundwork for proteome-scale endogenous tagging. To this end, we first show that low multiplicity of infection (MOI) lentiviral delivery of pegRNAs is compatible with efficient gene tagging that increases over time in cells with continuous expression of prime editing and split fluorescent protein systems. We next develop a two-step cloning strategy for the construction of pooled pegRNA libraries and build a large-scale library to exhaustively tag 60 endogenous proteins covering diverse localization patterns, while exploring a wide range of pegRNA parameters not previously tested in pooled combinations for their association with peptide knock-in efficiency across native genomic loci. Our analysis revealed both genomic and pegRNA features influencing tagging efficiency including chromatin state, strand nick location and pegRNA extension composition. Next, we integrated multiple features with small effects into a predictive model for pegRNA-mediated gene tagging. Lastly, we coupled pooled tagging with in-situ pegRNA sequencing and deep learning image analysis to cluster tagged cells based on observed protein localizations, establishing an end-to-end workflow for the scalable exploration of subcellular proteome organization.

## Results

### Low MOI pegRNA transduction enables efficient protein tagging over time

In practice, adapting perturbation or genome editing methods to a pooled high-throughput format targeting different genomic loci in mammalian cells requires that high perturbation or editing efficiency can be achieved with low MOI lentiviral transduction. To test prime editing mediated tagging efficiency with low MOI delivery of pegRNAs, we first designed a piggybac vector for the delivery of the prime editor (PE2) with both antibiotic and fluorescent based selection, to be used together with the mNG3K_1-10_ component of a split fluorescence protein system^17,18^ (Fig. 1A). We then engineered cell lines stably expressing both components using the following procedure: cells were first transfected with the piggybac donor and transposase followed by blasticidin selection. Selected cells were then transduced at high MOI with lentiviral particles containing mNG3K_1-10_ and transiently transfected with an exogenous CLTA gene fused to mNG2_11_. We then selected clonal cell lines for expression of both components using a near-infrared FP (emiRFP670^19^) and green fluorescence as indicators for expression of PE2 and mNG3K_1-10_ respectively (Fig. S1).

**Figure 1.**
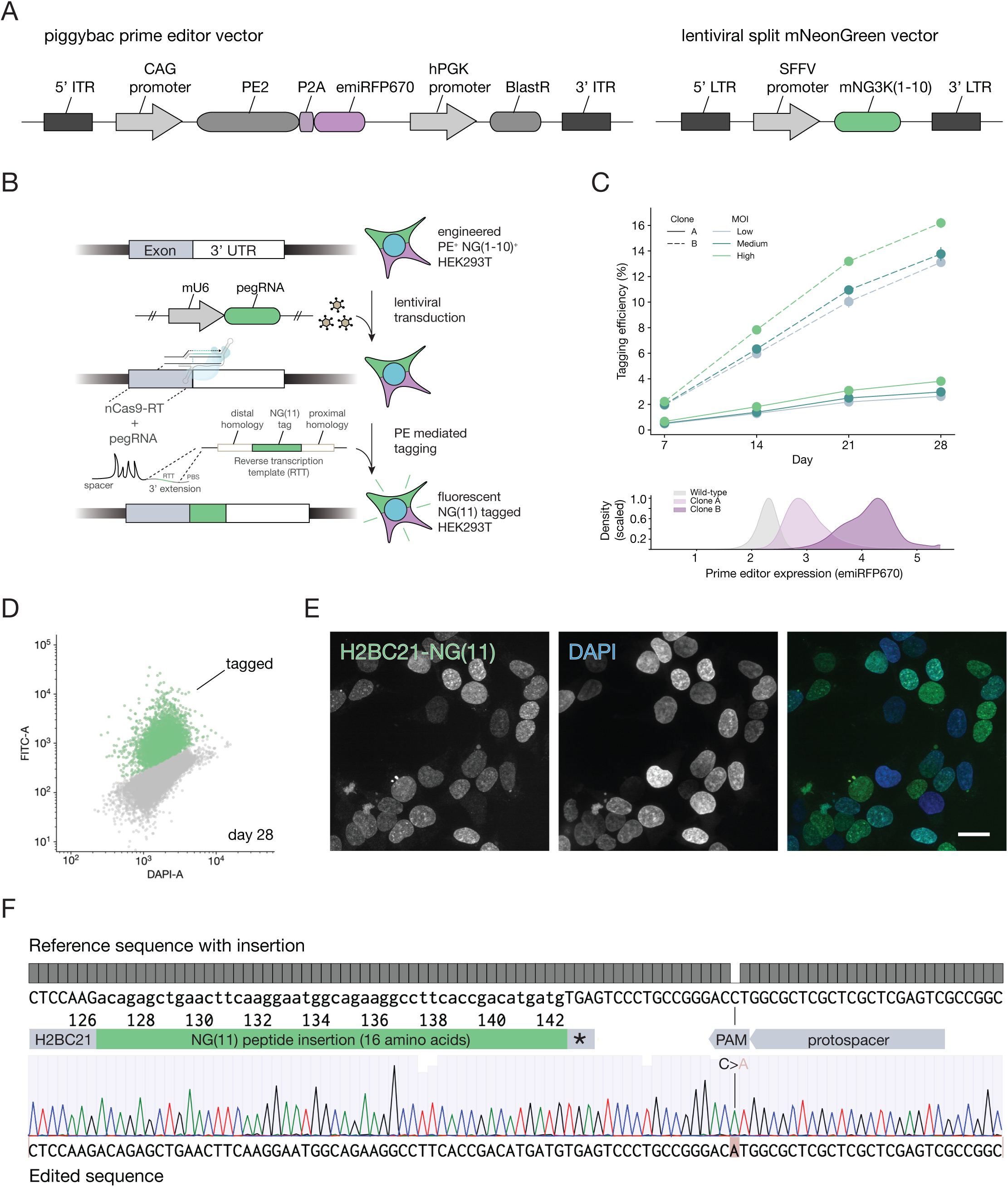
Stable expression of prime editing components enables endogenous tagging with increased efficiency over time after pegRNA delivery at low multiplicity of infection. (A) Vector design for piggybac prime editor and lentiviral split mNG(1-10) reporter (addgene #157993). (B) Workflow for prime editing mediated endogenous protein tagging in an engineered prime editing and split mNeonGreen reporter HEK293T line. (C) Tagging efficiency over time for two clonal lines expressing PE2 at different expression levels and delivery of an mNG(11) peptide insertion encoding pegRNA targeting the C terminus of H2BC21 at different MOIs. Error bars denote the standard deviation of n = 3 individual pegRNA transductions for each timepoint and condition. (D) Flow cytometry density plot at day 28 post lentiviral transduction of a pegRNA targeting the C-terminus of H2BC21 for mNG(11) tagging (E) 60X confocal images of mNG(11) tagged H2BC21 in post-sort HEK293Ts. Scale bar = 20 um. (G) Sanger sequencing trace for an mNG(11) tagged allele with respect to a wild-type reference allele from sorted tagged cells.

After establishing stable cell lines with both components, tagging endogenous proteins can follow from delivery of pegRNAs encoding the mNG2_11_ peptide targeted to either the N or C terminus of the target protein followed by self-complementation with the stably expressed mNG3K_1-10_ (Fig. 1B). To test the efficiency of tagging using this approach, we produced lentiviral particles containing a pegRNA to endogenously tag H2BC21 gene at the C-terminus and transduced our engineered clonal lines at different MOIs. We tested two clonal lines, with high and low PE2 expression and measured tagging efficiency using flow cytometry at different time points (Figure 1C,D). In both clones we observed an increase in tagging efficiency over time with the PE2 high-expressing clone achieving more than 10% tagging efficiency at day 28. We validated tag insertion precision by both microscopy of sorted polyclonal cells (Figure 1E) and sequencing of the target site (Figure 1F). Thus, using clonal lines selected for high PE2 and mNG3K_1-10_ expression we are able to achieve high efficiency endogenous tagging with low MOI pegRNA delivery, providing the basis for endogenous protein tagging in a pooled format.

### Design and construction of a large scale pooled pegRNA library for exploring design parameters associated with tagging efficiency

We next set out to test the efficiency of pooled protein tagging across many genomic loci and explore genomic and pegRNA features associated with tagging efficiency. We chose 60 genes with validated and diverse N and C terminus tagged localization patterns (30 for each terminus) using data from the OpenCell database^1^ and selected 4 targeting spacers for each (Figure 2A). For each gene, spacers were chosen by dividing the flanking sequence around the desired insertion site into 15 bp bins and choosing the spacer with lowest off-target CFD score when possible^20^. For each of these targeting spacers we designed 72 pegRNAs exploring various parameters including primer binding site (PBS) length, distance between the nick and edit, tag insertion size and RTT homology length distal to the nick site (Fig. 2B). A resulting library of 17,280 pegRNAs was divided into three sublibraries corresponding to three tag insertion sizes of 48, 60 and 78 nt. We also added a unique 12 bp barcode downstream of the pegRNA 3’ extension for identification of each construct by bulk amplicon sequencing and in-situ barcode sequencing. Each pegRNA was designed with a twin containing synonymous codon mutations in the tag insertion template to serve as internal replicates. We also included 240 non-targeting control pegRNAs in each insertion length sublibrary. To construct the pooled pegRNA plasmid libraries, we developed a two-step cloning procedure in which the spacer and 3’ extension were first synthesized together separated by two type II restriction enzyme sites and cloned into a CROPseq vector we modified to accept pegRNA libraries and retain compatibility with single cell and optical sequencing approaches. In the second step, the constant sgRNA scaffold was inserted through another large-scale cloning reaction (Fig. 2B,S2 and methods).

**Figure 2.**
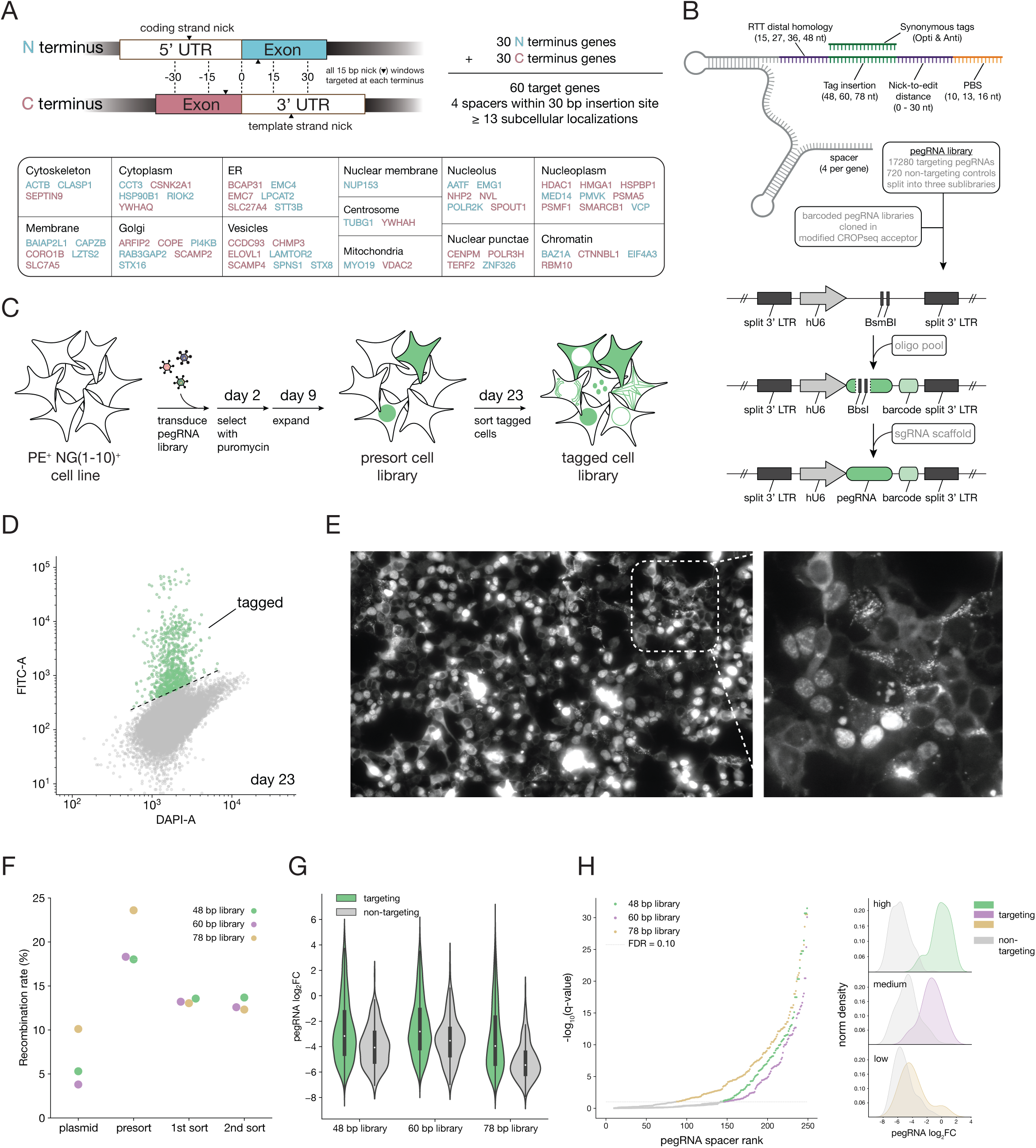
Construction of pooled pegRNA libraries for N and C terminus tagging of multiple genes while exploring a range of pegRNA design parameters. (A) Selection of genes covering diverse subcellular localization patterns, noting their terminus tagging and nicking locations. (B) Combinations of pegRNA design parameters explored for each spacer sequence. Tagging libraries are cloned in two consecutive large scale cloning steps starting with a pegRNA oligo pool. (C) Illustration depicting the generation of a pooled tagged cell library. (D) FACS 2D density plot at day 23 post pegRNA library lentiviral transduction for the 60 bp tag library. (E) Fluorescence image of a pooled tagged cell library showing diverse localization patterns, suggesting a different protein is tagged in each cell. (F) Spacer barcode recombination rates at different stages of library construction. (G) Violin plots showing targeting and non-targeting control distributions of pegRNA fold changes (log_2_FC for each double sorted sublibrary. As expected most pegRNAs are depleted in sorted cell populations with a long tail of active pegRNAs observed primarily in the targeting group. (H) Spacer ranking by FDR corrected p-values for a one-sided KS test comparing the distribution of pegRNAs for each spacer with the distribution of non targeting pegRNAs within each sublibrary. 108, 105 and 163 out of 240 spacers are considered active at a false discovery rate of 10% for sublibrary 48 bp, 60 bp and 78 bp respectively. 41, 48 and 56 out of 60 genes have at least one spacer considered active at a false discovery rate of 10% for sublibrary 48 bp, 60 bp and 78 bp respectively. All genes had at least one active pegRNA spacer across the three sublibraries. Right panel shows example pegRNA fold change (log_2_FC) distributions of spacers with high, medium and low tagging efficiencies.

To construct the pooled tagged cell library, similar to standard CRISPR screening methods, we produced lentiviral particles from the pooled pegRNA plasmid library followed by low MOI (<0.1) transduction and double sorting to isolate the polyclonal population of cells with successful tagging (Fig. 2C). Successful tagging was apparent from FACS analysis during library sorting (Fig. 2D) and from fluorescence imaging that revealed a mix of localization patterns suggesting that a different protein had been endogenously tagged in different cells (Fig. 2E). As our pooled pegRNA library contains variable regions (spacer, 3’ extension and barcode) separated by constant sequence, it is prone to recombination during PCR and template switching during lentiviral production^21,22^. To mitigate that risk, we minimized the distance between the pegRNA and barcode sequences, used a low number of PCR cycles and quantified barcode misidentification using long cycle paired end sequencing at different steps of plasmid and cell library construction (Fig. S3). We found recombination rates similar to previous studies with decreased rates in the sorted cell samples, likely due to mismatch between the spacer sequence and 3’ pegRNA extension resulting in failed tagging and no fluorescence signal (Fig. 2F). Bulk analysis of the read count distributions of the three sublibraries across cloning and cell library construction stages showed sufficient coverage in plasmid library and initial cell library transduction (Fig. S4) followed by increased skew after sorting for tagged cell libraries as expected (Fig. S5,6).

To gain insight into pegRNA features that are associated with increased likelihood of successful tag insertion, we first calculated tagging efficiency as the log2 fold ratio of counts between double sorted and presorted cell libraries for each pegRNA (Table S1). As each spacer sequence was incorporated in many pegRNAs with varying extension parameters, we could compare tagging efficiency distributions of pegRNAs by targeting and non-targeting spacers, which revealed increased efficiency for the former, yet with high variability (Fig 2G). To gain a rough estimate of the number of spacers that resulted in successful gene tagging, we compared the distribution of tagging efficiency for each spacer to the distribution of pegRNAs with non-targeting control spacers and calculated an FDR corrected q-value based on a one sided Kolmogorov–Smirnov statistic (Fig. 2H). Based on this analysis, 139, 125 and 135 spacers out of 240 in the 48, 60 and 78 bp insertion size sublibraries respectively were tagging at a 10% false discovery rate. All genes had at least one spacer that resulted in successful tagging across all three sublibraries (Fig. 2H). This suggests that a majority of the genes we set out to tag can be tagged using low MOI lentiviral pegRNA delivery. These numbers can vary as a function of cell coverage at transduction and sorting, yet it suggests that prime editing is sufficiently efficient across many genes to facilitate large-scale tagging efforts.

### Tagging efficiency varies significantly across genes, epigenetic states, termini and strands

We next set out to examine the contributions of genomic and pegRNA features to tagging efficiency. Examining tagging efficiency distributions of individual genes, ranked by average, reveals that while the gene identity plays an important role in determining tagging efficiency, there is also significant variability, suggesting contributions from spacer choice and pegRNA extension features as well (Fig. 3A). Comparison of the average tagging efficiency for each gene between sublibraries with different insertion sizes revealed the effect of the gene is highly reproducible, suggesting that it is determined by genomic features and can be used in future library design to reduce the skew in representation of tagged proteins (Fig. 3B,S7). Target gene expression levels were somewhat correlated with tagging efficiency, with the fluorescence measurements of the tagged protein, taken from arrayed measurements^1^ showing the highest correlation (Fig. 3C). Interestingly, we found that tagging was more efficient for N terminally tagged genes, which might be due to chromatin state or interactions with gene expression regulation machinery (Fig. 3D).

**Figure 3.**
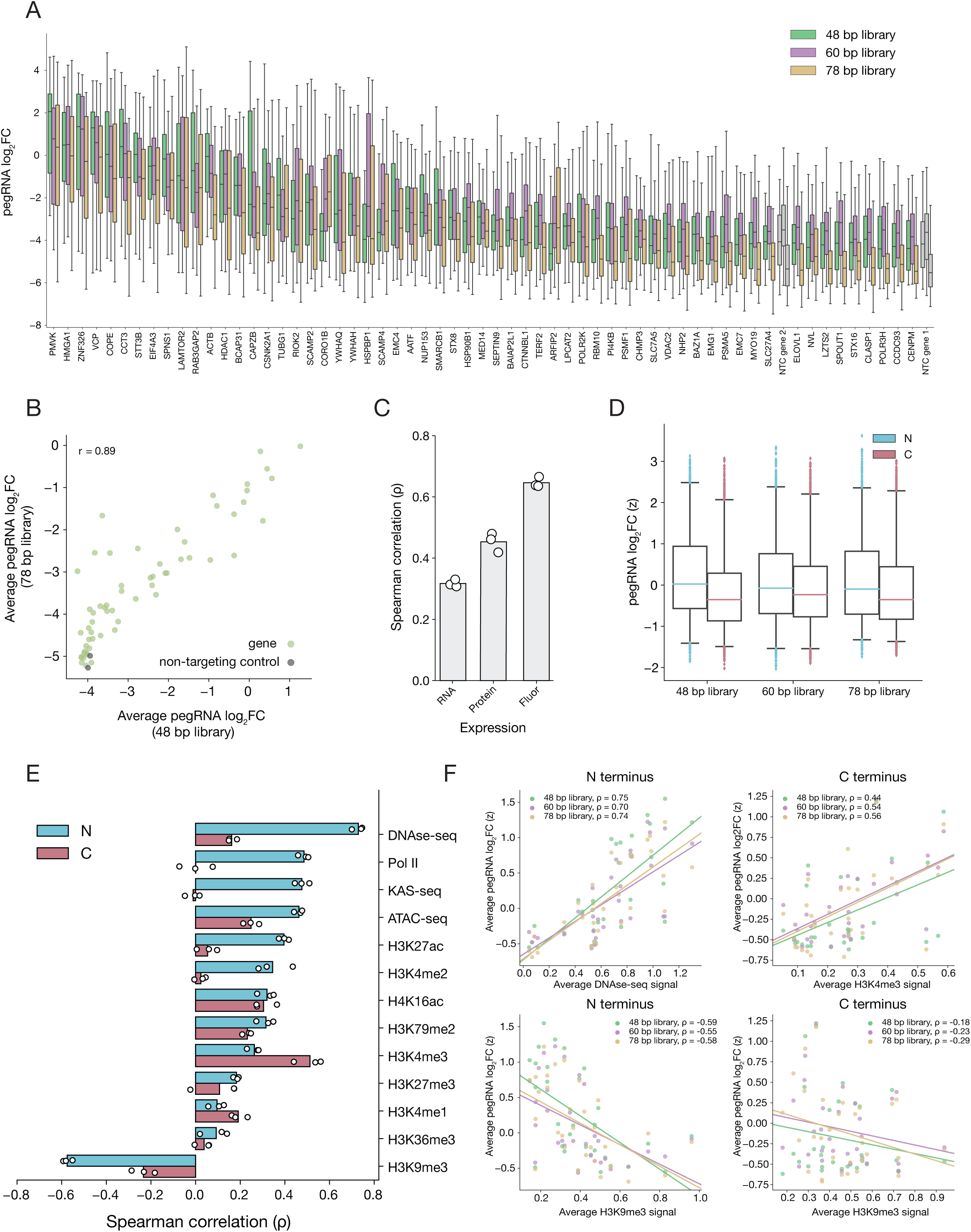
Tagging efficiency varies across genomic loci and chromatin states. (A) Distributions of pegRNA tagging efficiencies for each gene ranked by average gene effect (mean pegRNA log_2_FC). Whiskers indicate 2.5th and 97.5th percentiles. (B) Comparison of the average gene effect between the 48 bp and 78 bp tag sublibraries. Pearson r correlation is noted. (C) Correlation between the average gene tagging efficiency and measurements of mRNA, protein and fluorescence expression. Highest correlation is observed when tagging efficiency is compared to fluorescence expression by FACS of the same genes tagged in an array format (from OpenCell). (D) Distribution of tagging efficiency comparing tagging at the N and C termini. Whiskers indicate 2.5th and 97.5th percentiles. (E) Correlation between tagging efficiency and different chromatin features sorted in descending order by the correlation for N terminus tagged genes. (F) Example correlations of both the N and C termini with top positive and top negative associated chromatin features.

To systematically evaluate if epigenetic markers are associated with differences in tagging efficiency, we curated publicly available datasets ^23–33^ for HEK293T and HEK293 (when HEK293T data was not available) and calculated the average signal across a 2kb window centered on the start codon or stop codon insertion points of N terminus and C terminus targeted genes respectively. We then calculated the correlation between the average gene log2FC tagging efficiency and the average feature signal and sorted the features by decreasing correlation with respect to the N terminus (Figure 3E). We found that in general the average tagging enrichment of C terminus genes did not correlate as strongly with these genomic features as the N terminus did. Looking through each genomic feature association, we found that the N terminus genes had a spearman correlation of ρ ≈ 0.7 for the corresponding signals in the DNAse-seq dataset, suggesting an open chromatin state is a strong indicator of the expected average tagging efficiency at the N terminus, while at the C terminus this correlation was lower but still positive in direction. In general, open chromatin features such as ATACseq and transcriptional activity features such as KAS-seq also had moderate positive correlations, ρ ≈ 0.5, supporting the importance of chromatin accessibility to the prime editor complex. We also found that euchromatin associated histone modifications had overall positive correlations with average tagging enrichment, while heterochromatin associated H3K9me3 marks, known to be associated with transcriptional repression, had a negative correlation. Interestingly, we noticed that the H3K4me3 mark had the strongest positive correlation for the C terminus but not for the N terminus (Figure 3E). Altogether, this analysis emphasizes the importance of chromatin state as a key feature influencing prime editing insertion efficiency, which might also explain the difference in tagging efficiency between the N and C terminus targeted genes. These findings are in line with a recent study^34^ that assessed the influence of chromatin features on prime editing mediated 3 bp insertions within randomly integrated piggybac target sites, suggesting the prime editability of a genomic location is less dependent on insertion length and more coarsely tuned by epigenetic states of the target.

Next, given the strong association of tagging efficiency with transcriptional state, we explored potential differences when tagging is initiated by nicking the transcriptional coding strand or template strand. In both N and C terminus insertions, nicking upstream of the tag insertion site would require nicking the coding strand and using a pegRNA-encoded tag that is reverse complement, while nicking downstream of the insertion site would require nicking the template strand and using a pegRNA-encoded tag that has the precise coding tag sequence (Fig. 4A). We found that in both N and C termini, nicking the coding strand resulted in higher tagging efficiency (Fig. 4B). Gene tagging by nicking the coding or template strand required the use of two distinct insertion template sequences within the pegRNA extension. We found these had different abundances within the library, which is likely due to differential oligo synthesis or PCR amplification (Fig. S8A). We ensured that the differences in tagging efficiency between the strands was independent of the initial pegRNA abundances in the sublibraries (Fig. 4C) and was observed at the level of individual genes. Indeed, most genes showed reduced tagging efficiency when comparing the average fold change of pegRNAs programmed for a template strand nick compared to a coding strand nick (Fig. 4D). This effect was also independent of predicted spacer efficacy score, which can also be affected by differences in nucleotide composition between the two strands (Fig. S8B). Binning pegRNAs by nick position revealed a similar difference between upstream and downstream nicking locations (Fig. 4E), in agreement with our strand analysis (Fig. 4B). We also observed increased tagging efficiency as the nick position was closer to the tag insertion site both on average (Fig. 4E,F) and when examining individual genes (Fig. 4G). Analysis of the strictly standardized mean difference (SSMD) between average fold changes of pegRNAs targeting the coding and template strand for each gene further suggested that most genes had a coding strand bias associated with positive correlations against RNA expression that increased slightly with the length of the insertion (Fig. 4H) and greater than against protein expression measurements (Fig. 4I).

**Figure 4.**
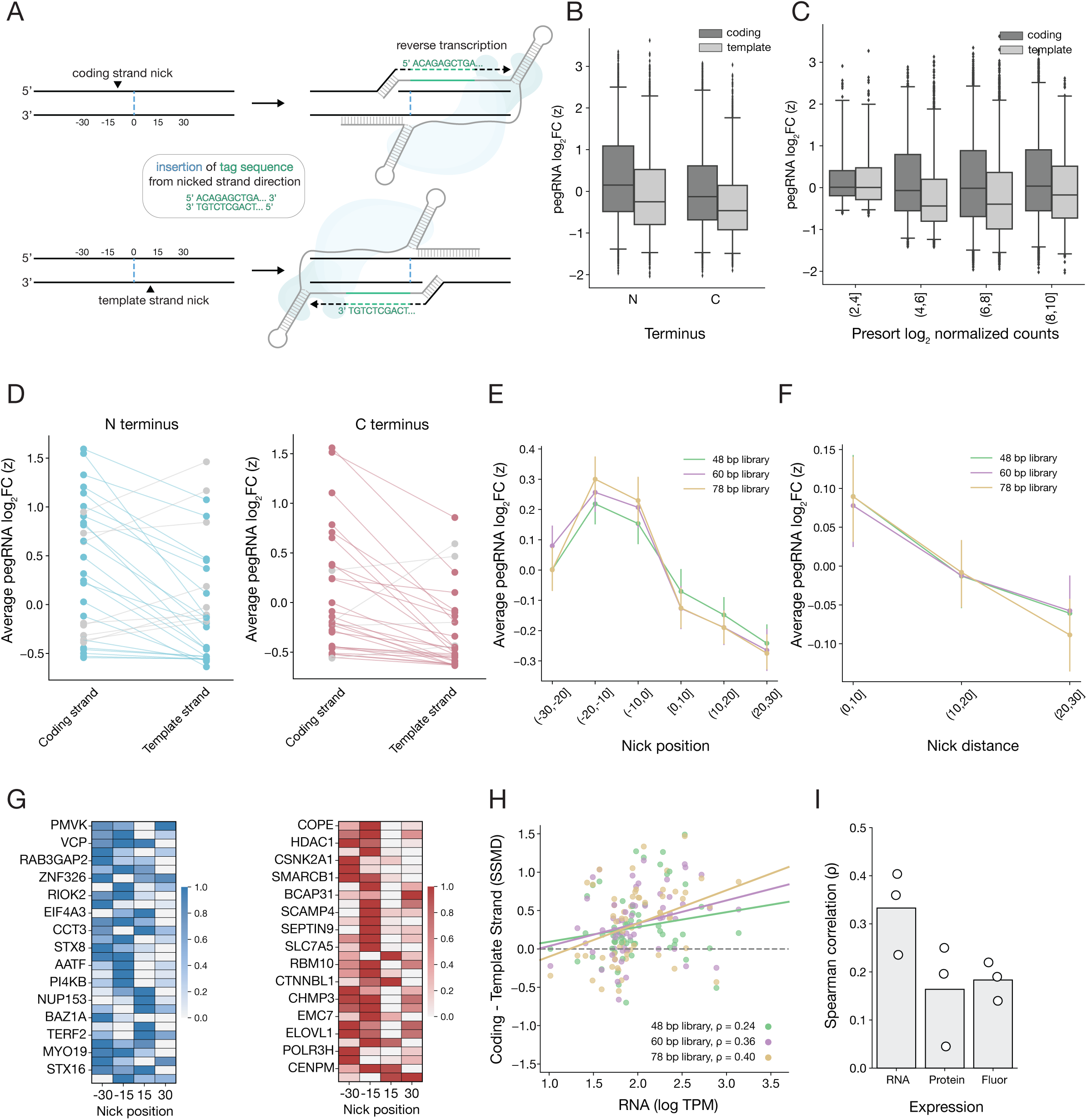
Nicking and insertion through the coding strand is associated with higher tagging efficiencies. (A) Illustration of how strand nicking location determines pegRNA insertion template design. (B) Box plots of tagging efficiency showing increased rates nicking the coding strand across N and C termini. Whiskers indicate 2.5th and 97.5th percentiles. (C) Box plots of tagging efficiency showing increased rates when nicking the coding strand. pegRNAs are binned by presort day 23 abundance across sublibraries. Whiskers indicate 2.5th and 97.5th percentiles. (D) Reduced average tagging efficiency when nicking is performed on the template strand for each individual gene. (E) Average tagging efficiency as a function of nick position relative to tag insertion site. Error bars indicate 95% confidence intervals. (F) Average tagging efficiency as a function of nick distance to tag insertion site. Error bars indicate 95% confidence intervals. (G) Heatmap showing average tagging efficiency min-max normalized for each gene and binned by nick position. (H) Strictly standardized mean difference (SSMD) of average tagging efficiencies for pegRNAs nicking the coding strand and template strand for each gene.Shows a positive correlation with RNA expression that increases with insertion size. (I) Difference between tagging efficiency at the two strands correlates more with RNA expression than protein measurements.

### Systematic analysis of pegRNA features associated with tagging efficiency

We next explored different pegRNA features for association with tagging efficiency. We first examined the spacer on-target score^35^ and found higher scores to be associated with higher tagging efficiency (Fig. 5A). Closer examination of this relationship suggested that an on-target score > -0.5 may be necessary for an efficient tagging pegRNA but not sufficient, as the pegRNA spacers cover both low and high tagging enrichments in this on-target score range (Fig. 5B). The PBS is another component of the pegRNA for which sequence composition may have an effect on activity. Thus we focused on three PBS-associated features: length, GC content and melting temperature. We found that the tested lengths of 10, 13 and 16 nt did not differ significantly in their average tagging efficiency (Fig. 5C) but noticed instead that GC content and melting temperature had clearer value ranges that on average led to higher tagging efficiencies (Fig. 5D) with high GC content and melting temperatures being favorable. These PBS GC content and melting temperature trends agree with previous literature that explored pegRNA features influencing the efficiency of point mutations and smaller insertions^10–12^, suggesting that favorable PBS sequences are independent of the type of prime edit.

**Figure 5.**
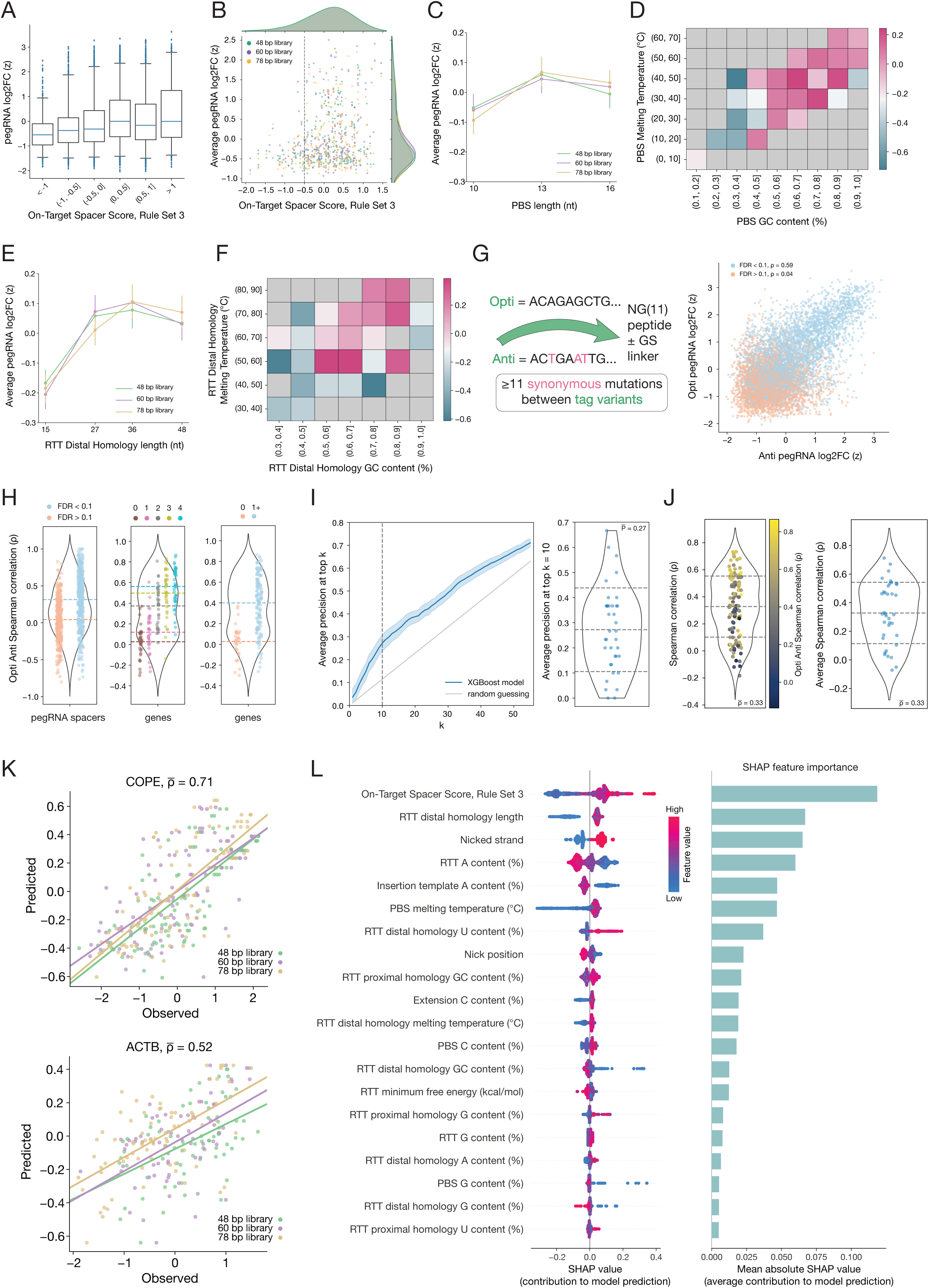
pegRNA design features have small effects on tagging efficiency that can be integrated into a predictive model. (A) Tagging efficiency distributions when pegRNAs are binned by on target spacer score (RS3). Whiskers indicate 2.5th and 97.5th percentiles. (B) Comparison of average tagging efficiency for each spacer against on target spacer score (RS3) suggests a spacer score threshold necessary but not sufficient for efficient tagging. (C) Tested PBS lengths have similar tagging efficiencies. Error bars indicate 95% confidence intervals. (D) Heatmap comparing tagging efficiency for different value ranges of PBS GC content % and melting temperature °C. (E) Larger RTT homology sequences distal to nick site result in higher average tagging efficiencies. Error bars indicate 95% confidence intervals. (F) Heatmap comparing tagging efficiency for different value ranges of RTT GC content % and melting temperature °C. (G) Comparison of tagging efficiency for pegRNA pairs with synonymous tag insertion template sequences. Left panel illustrates the pegRNA pair design with synonymous mutations that generate different tags with the same resulting amino acid sequence. Right panel shows a scatter plot of pair values colored by FDR compared to non-targeting spacers (Figure 2H). (H) Correlation between Opti/Anti pegRNA pairs to evaluate reproducibility of pegRNA design feature effects on tagging efficiency. Left most panel shows this correlation at the level of individual pegRNA spacers, separating spacers that passed FDR for tagging from those that did not. Middle panel at the level of genes separated based on the number of spacers that passed FDR. Right panel separates genes based on at least one spacer that passed FDR. Dashed lines indicate mean. (I) Precision (y-axis) of predicting the top K most active pegRNAs (x-axis) using XGBoost model compared to random guessing, showing that the model is able to predict active pegRNAs with the tested features. Right panel shows precision for individual genes revealing variability in capture of most active pegRNAs across genes. Center dashed line indicates mean and outer dashed lines indicate one standard deviation. (J) Correlation between predicted and observed tagging efficiencies for each gene demonstrating variability in predictive ranking of pegRNAs across genes. In left panel genes are divided by sublibraries and in right panel each gene is averaged across sublibraries. Center dashed line indicates mean and outer dashed lines indicate one standard deviation. (K) SHAP analysis of feature contributions to XGBoost model predictions. Top 20 features shown are sorted based on feature importance (mean absolute SHAP value). Nicked strand is a categorical feature, red indicates coding strand nick and blue indicates template strand nick. (L) Examples of predicted and observed tagging efficiencies for two genes with high and intermediate correlations.

The next pegRNA region we specifically examined was the RTT homology distal to the nick site, focusing on length, GC content and melting temperature. We found that short homology lengths of 15 bp tended to have lower tagging efficiencies on average than longer homology lengths ≥ 27 bp (Fig. 5E). We also found that high GC content and high melting temperatures tended to result in higher tagging efficiencies following a similar trend as for the PBS (Fig. 5F). Altogether, this suggests that increased structural stability of the 3’ extension may be an important factor driving tagging efficiency. Lastly, to assess the reproducibility of our measurements, we compared the tagging efficiencies of pegRNA pairs that were designed with synonymous tag mutations as internal replicates for pegRNA design. We found that for pegRNA pairs whose spacers passed FDR, likely associated with true protein tagging, these measurements correlated well, suggesting that indeed we are able to capture pegRNA design parameters associated with increased efficiency (Fig. 5G).

### Features with small effects can be integrated into a predictive model of more efficient tagging pegRNAs

As we found multiple pegRNA features with small effects on tagging efficiency, we asked if integrating them into a predictive model would result in larger increases in efficiency that could be used for the design of improved pegRNAs for gene tagging. Towards that goal, we used gene normalized pegRNA efficiencies for 37 genes that passed FDR (Fig. 2H) such that genes with larger dynamic range of both successful and unsuccessful pegRNAs were included. We trained a gradient boosted trees regression model (XGBoost)^36^ on an initial set of 63 pegRNA features, including on-target spacer score (Rule Set 3)^35^, nicked strand, nick position relative to insertion point, nick distance from insertion point, PAM disruption, first nucleotide of the 3’ extension as well as length, nucleotide content, GC content, melting temperature and mean free energy^37^ for different regions of the pegRNA. The model was evaluated through nested cross-validation with a leave-one-gene-out (LOGO) strategy for testing and grouped k-fold cross-validation (k=6) for hyperparameter tuning within inner folds. Performance was evaluated using the average spearman correlation between predicted and observed tagging efficiencies across genes. To rank feature importance, we performed Shapley additive explanations (SHAP) analysis^38,39^ and removed features that did not contribute to model predictions. This feature selection and model building process was repeated iteratively, converging on a set of 37 features used for the final model (see methods).

To evaluate the predictive value of our model, we used the average precision at top k pegRNAs and the correlation between observed and predicted efficiencies for each gene when the pegRNAs of that gene were left out as the test set. We found an average top k precision of about 0.3 at k=10 (Fig. 5I, left panel) with variability in predictive value across genes (Fig. 5I, right panel). Similar results were observed when we used the spearman correlation between predicted and observed efficiencies to evaluate model ranking performance, where the model could properly rank pegRNAs for some genes and less for others (Fig. 5J,K). Notably, model performance was higher for genes that showed a high correlation between opti- and anti-pegRNA pairs (Fig. 5J, S10), suggesting that pegRNA measurement error plays a role in the variable predictive value across genes. Final ranking of feature importance by SHAP analysis revealed features associated with all parts of the pegRNA, including spacer on-target score, nicked strand, RTT and PBS compositions, contributing to increased tagging efficiency (Fig. 5L). In summary, our analysis demonstrates the integration of multiple features into predictive models for increased tagging efficiency and emphasizes the importance of measuring pegRNA mediated editing across multiple genomic loci.

### Coupling pooled tagging with in-situ sequencing provides a snapshot of subcellular protein organization

Imaging-based phenotyping of pooled tagged libraries offers the opportunity to investigate localization patterns of many proteins in parallel and provide a birds-eye view of subcellular organization. Motivated by this idea, we cloned the pegRNA libraries in a modified CROPseq vector so that both the pegRNA and downstream barcode will be transcribed into a polyadenylated transcript^40^ that can be detected by *in-situ* Sequencing-By-Synthesis (SBS)^41,42^. In order to generate a large imaging based dataset of single cell localization patterns for each pegRNA within the pool, we used high-throughput automated widefield fluorescence microscopy. First, localization patterns of the tagged proteins were phenotyped at a higher resolution (40x) together with a phalloidin stain for cellular actin-based segmentation and DAPI for nuclear segmentation. Next, we proceeded with an in situ sequencing workflow to identify the pegRNA in each cell^41,42^ at 10x with an additional pre-RT fixation step using amine-modified RT primers to improve retention of targeted RNA transcripts and cDNA products in situ^43^. We tested the use of two different padlock sequences, one to capture the pegRNA spacer and a second to capture the barcode. Individual cells were mapped between the two imaging magnifications such that the identity of each tagged protein could be determined (see methods). We then used the actin-based segmentation mask to crop up to 1500 cells for each pegRNA spacer, resulting in a single cell imaging dataset for each tagged protein (Fig. 6A).

**Figure 6.**
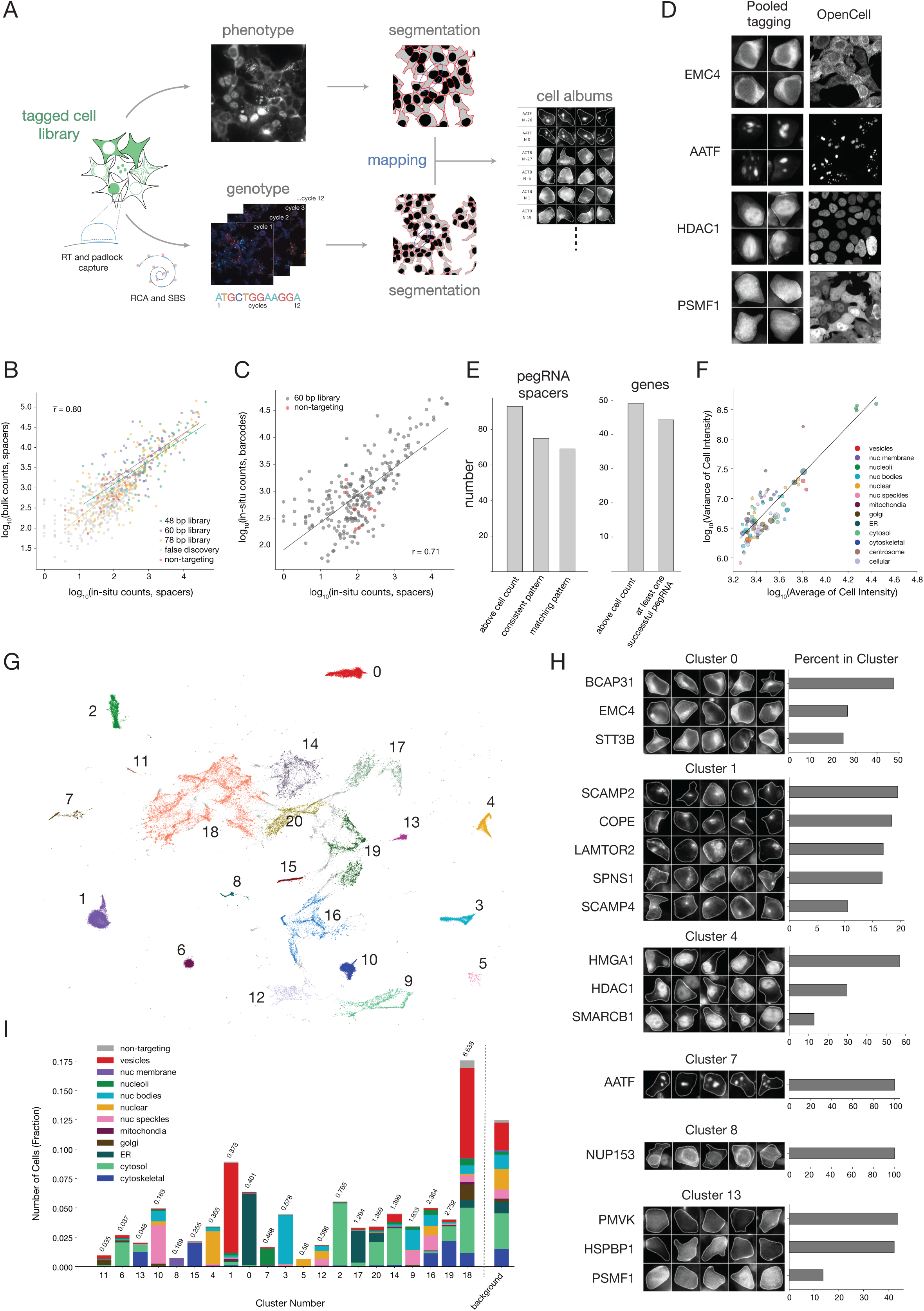
Coupling pooled tagging with in-situ sequencing enables the capture of subcellular localizations of proteins in parallel. (A) Illustration of the data acquisition pipeline for generating cell albums of individual tagged proteins out of a pooled tagged cell library. (B) Comparison of in situ sequencing cell counts with bulk sequencing normalized read counts for pegRNA spacers in each sublibrary. Pearson r correlation is noted. (C) Comparison of in situ sequencing cell counts using two different padlock probes capturing pegRNA spacers and downstream barcodes respectively for the 60 bp tag insertion sublibrary. Pearson r correlation is noted. (D) Example of individual tagged proteins in comparison with images from OpenCell database where the same protein was tagged in an arrayed format. (E) Manually evaluated statistics on tagging success based on evaluation of cell albums. (F) Correlation of average tagged cell fluorescence intensity and cell-to-cell variability in intensity showing that variance in protein expression scales with protein expression. (G) Two dimensional UMAP embedding of the latent parameters inferred by the trained cytoself neural network followed by HDBSCAN clustering. (H) Example cell albums for clusters with the top represented genes in each cluster and the percentage of cells within the cluster for each tagged gene based on in situ sequencing. (I) Composition of localization patterns within each cluster as determined by HDBSCAN. Expected localization pattern for each protein is based on condensed OpenCell annotations.

In-situ cell counts for all three sublibraries correlated well with normalized read counts from bulk amplicon sequencing (Fig. 6B). Similarly cell counts using either of the two padlock sequences were well correlated, with low cell counts for non-targeting pegRNAs likely representing cells with more than one pegRNA lentiviral integration (Fig. 6C). We next took a conservative cell count cutoff using the most highly represented NTC, which corresponded to 513 cells, and generated cell albums for each pegRNA spacer passing the cutoff (Fig. S11-14). Manual investigation of the resulting spacer albums revealed consistent localization patterns for the vast majority of spacers, which reassuringly, matched external arrayed tagging datasets^1^ (Fig. 6D,E and Fig. S11-14). As our datasets measure protein localization and fluorescence in hundreds of cells for each tagged protein, it can also provide valuable information on the variability in these measurements across single cells. In line with this, we calculated the average fluorescence intensity and variability for each tag and found that the variability in protein abundance scaled with average abundance as was previously observed^44^ (Fig. 6F).

We next tested if protein localization patterns can be clustered in an unbiased manner, to provide a birds-eye view of cellular organization. We used our dataset of all the spacer albums that passed the NTC cell count cutoff to train a self-supervised Variational AutoEncoder (VAE) previously designed to analyze cellular protein localization images^45^. Using the trained VAE we inferred the latent space representation of each tagged cell which was then flattened and embedded onto a 2D UMAP (Fig. 6G). To identify distinct clusters we performed unsupervised clustering via HDBSCAN. This resulted in dense clusters at the periphery which showed highly consistent localization patterns for genes enriched within each cluster (Fig. 6G,H and Fig. S15). To enable systematic characterization of each individual cluster, we ranked them by the normalized K-Means cluster score that captures the spread of each cluster (see methods). We then plotted the composition of expected localization patterns in each cluster determined using OpenCell annotations (Fig. 6I, Table S2). We found that for diffuse clusters (K-Means score > 2) several annotated localization patterns can be observed (clusters 16, 18 and 19), likely due to cells with dim and unclear fluorescence signals. Conversely, the majority of clusters with K-Means Score < 2 consisted of cells with a similar expected pattern with most inconsistencies being associated with closely related patterns (e.g. nuclear speckles and nuclear bodies in cluster 10). Altogether, this shows that pooled protein tagging via prime editing can be used to gain insight into cellular organization of many proteins in parallel, enabling scalable studies of subcellular dynamics in response to genetic changes and external stimuli across cell types.

## Discussion

In this study, we took advantage of pegRNAs ability to encode both target nick sites and insertion templates within single lentiviral vectors to develop scalable pooled protein tagging for multiplexed imaging of subcellular protein localization. We designed prime editing insertion libraries containing 17,280 pegRNAs targeting 60 genes to explore genomic and pegRNA features influencing endogenous tagging success. Most of the genes we attempted to tag were successful and resulted in localization patterns that matched expectations, suggesting that prime editing can be used to effectively tag genes across diverse genomic loci for exploration of proteome organization. Our work establishes an end-to-end workflow for the generation of pooled tagged cell libraries and their analysis by combining high-throughput imaging with in-situ barcode sequencing and deep learning based image analysis.

While we were able to tag the majority of intended genes, we observed large differences in efficiencies driven by both gene and pegRNA features. We found that tagging efficiency varied significantly between genes and correlated with epigenetic features associated with open chromatin and active transcription. We also found multiple features associated with the choice of pegRNA that increased efficiency including on-target spacer score, nicking location and nucleotide composition of PBS and RTT distal homology regions. As each feature had a small effect on tagging efficiency, we integrated multiple features into a predictive model that enables in silico ranking of pegRNA designs for a given gene – to constrain the search space of active tagging pegRNAs for future libraries. Model performance varied across genes, likely due to differences in coverage that affected the accuracy of the measurements (Fig. S9). Pooled protein tagging offers a distinct strategy for testing genomic and pegRNA features at scale to model prime editing efficiency and offers complementary insights to self-targeting random integration^10–12^ and single genomic loci approaches^13^.

One pegRNA feature found to be strongly associated with higher tagging efficiency was the choice of nicked strand with insertions through the coding strand resulting in higher efficiency. This has not been observed in previous studies^10–13^, possibly because it only affects larger insertions, which previous work did not test systematically across both strands. Indeed, we find an increased effect with longer insertion lengths (Fig. 4H). A potential explanation for the coding strand bias in tagging enrichment is that Cas9 indel activity has been shown to be higher when the sgRNA binds the template strand due to RNA Pol II removing Cas9 from the target DNA during transcription, allowing more efficient access of host DNA repair factors for resolution of the double strand break^46^. Another potential explanation is that transcription-coupled nucleotide excision repair (TC-NER) may be a pathway involved in resolving prime editing mediated insertions in at least the 48 bp to 78 bp lengths tested. The template strand is known to undergo rapid TC-NER, while the coding strand is more likely to be repaired through slower global-genome nucleotide excision repair (GG-NER)^47^. Hence, leading to higher insertion rates through the coding strand relative to the template strand.

In this work we used N and C terminus tagging, as it provided us with expected localization patterns that we could use to validate our results^1^. While N and C terminus tagging is most commonly used for both exogenous and endogenous protein fusions, gene termini can also encode for localization signals and unstructured regions that mediate protein-protein interactions. As prime editing is not limited to the gene termini, future work can use protein structural information and fold prediction of the effect of the fusion to tag proteins internally in a manner that might even be less disruptive than the termini. Indeed, in our previous work, we have shown that pooled internal tagging can be performed to identify non-disruptive tagging locations^6^. As both endogenous protein tagging and immunohistochemistry approaches can affect the resulting localization pattern, information from multiple experimental sources is required, including tagging proteins in more than one site. The scale of pooled protein tagging can be used to accommodate multiple tag sites per gene and increase the confidence in measurement of protein localization across cell types and conditions.

The development of scalable pooled tagging and imaging technologies, as presented here, will provide a valuable tool to study subcellular dynamics in a holistic manner. This could include systematic studies of changes in subcellular protein localization in response to environmental stimuli. As there are well-established stress-associated transient membraneless organelles such as stress granules and paraspeckles, it is likely that unbiased approaches will identify additional transient structures and rearrangements of subcellular organization that represent important aspects of cellular response to such perturbations. An additional application of pooled protein tagging could be to profile cellular disease models for subtle disease associated cellular phenotypes, which may not be apparent when using measurements of mRNA and protein abundance or localization of standard organelle markers. Such studies will also be driven by continuous advances in high-throughput imaging and analysis. In the current study, our imaging was limited to widefield microscopy using small cells that are not ideal for high resolution imaging of subcellular structures, yet as pegRNAs are delivered using lentiviral vectors, this approach can readily be expanded to other cell types. The use of pooled cell libraries with reduced skew will require less imaging per tagged protein supporting the generation of higher quality images using confocal microscopy or other higher resolution imaging methods. Such high quality imaging data, together with further advances in deep learning image analysis and more accurate and efficient in-situ sequencing^54^, will dramatically increase the granularity of localization pattern classification and even enable investigations into the variability in protein localizations across cells given the potential scales of these experiments.

Lastly, pooled epitope tagging can also be used for direct perturbation of proteins using nanobodies and degron systems, expanding the modalities of high-throughput DNA and RNA perturbation tools into the protein space. The development of next-generation tag systems will continue to unlock exciting applications of pooled tagging. Examples include new chemical biology tools and de novo designed peptide and peptide-binding protein systems^52,53^. Our work lays the foundation for future pooled tagging technology development and applications by demonstrating parallel and efficient tagging of many proteins across a wide range of genomic loci, providing rules for pegRNA design and establishing an experimental and computational framework for image-based analysis of endogenously tagged proteins – both to validate tagging accuracy and as a tool to study subcellular proteome organization at scale.

## Limitations of the study

A major challenge in the application of pooled tagging is that the resulting cell libraries are highly skewed in representation, due to the large differences in tagging efficiencies between genes. It is likely that increased cell and sequencing coverage, together with increased editing time, would have enabled the recovery of more genes, yet it would not have solved the increased skew within the library, which hinders imaging applications that are limited by surface area and high cost of in-situ sequencing reagents. Solving the skew can be done in several ways, including balancing the libraries by predicted or empirical measurements of efficiency. It is also likely that given the pace at which the prime editing field is advancing, more efficient systems will result in increased tagging that is less dependent on the epigenetic state of the target loci. Another challenge will be extending this pooled tagging approach beyond immortalized cell lines such as HEK293T used in this work to primary cells and induced pluripotent stem cells (iPSCs), where the efficiency of prime editing mediated tag insertions may be significantly lower. Nonetheless, continual improvements in prime editing efficiency through engineering and evolution of the prime editing system, better pegRNA design and modulation of endogenous DNA repair and chromatin states will help meet this challenge. Concurrently, increasing the size of insertions using single or paired pegRNAs^48–51^ while retaining scalability, or developing smaller fluorescence tags that obviate the need for a split tag system, will further expand the breadth of this pooled tagging approach.

## List of supplementary material items

Supplementary Figures S1-10

Fig. S11-14: Cell albums with representative cell images by pegRNA spacer

Fig. S15: Cell albums with representative cell images by cluster

Table S1: Normalized read counts and raw tagging efficiencies as determined by bulk sequencing

Table S2: Localization pattern annotations for each gene based on OpenCell and condensed annotations used in Figure 6F and 6I

## Acknowledgements

We would like to thank all members of the Shalem lab for support and discussions related to this project and Teodora Orendovici and Stephen Mahoney from the CHOP sequencing core. Majority of funding for this project came from NIH/NIGMS grant DP2-GM137416 awarded to O.S..

## Availability of data and materials

Raw pegRNA efficiencies are provided as a table in supplementary data including cell album images for pegRNA spacers. All plasmids used in this study will be deposited to Addgene upon paper acceptance. Code will be shared through GitHub.

## Methods

### Mammalian cell culture

Human Embryonic Kidney 293T cells (ATCC CRL-3216) were maintained in DMEM (Gibco 11995065) with 10% FBS, 1% NEAA (Gibco 11140076) and 1% Anti-Anti (Gibco 15240062). Cells were grown at 37°C with 5% CO2 to maintain physiological pH. Cells were tested for mycoplasma contamination on a monthly basis.

### Lentivirus production

Individual lentiviral particles were produced as follows: human 293Ts (ATCC CRL-3216) were plated such that they would be 75% confluent at the time of transfection in plates coated with 0.1% gelatin. Lentivirus for individual pegRNAs and pooled libraries was prepared in 15 cm plates by co-transfecting 293Ts with 13.25ug pMDLG, 7.2ug pMD2G, 5ug pRSV-Rev, 20ug individual pegRNA or plasmid library, 4 mL Opti-MEM, and 136 uL PEI per plate. For individual pegRNAs, media was replenished 6 to 6.5 hours post-transfection and 49 to 50 hours post-transfection the supernatant was collected and filtered through a 0.45μM filter. For pooled libraries, supernatant was collected 69 to 70 hours post–transfection. Supernatant was aliquoted and stored at -80C until use. For arrayed pegRNA transduction (Figure 1), lentivirus was thawed on ice and added to cells in suspension at multiple MOIs.

### Engineering HEK293T lines stably expressing prime editing and split fluorescent protein systems

Wild-type HEK293T cells were transfected with a piggybac transposon vector containing the PE2 system (Figure 1A) and piggybac transposase vector (SBI #PB210PA-1) followed by selection with blasticidin for two weeks. Polyclonal selected populations were then transduced with lentiviral particles containing mNG3K_1-10_ and transfected with a plasmid expressing CLTA fused to mNG2(11) at the N terminus to produce a fluorescent signal in cells expressing mNG3K_1-10_. Fluorescent Activated Cell sorting (FACS) was then used to isolate clonal lines that express both PE2 using emiRFP670 fluorescence and mNG3K_1-10_, using self-complemented mNeonGreen fluorescence. Clonal lines were characterized for having normal growth rate and the ability to endogenously tag H2BC21.

### pegRNA library design

We first chose 60 genes with validated localization patterns from the OpenCell database, 30 from each termini, spanning a diverse set of subcellular localizations including nucleoplasm, chromatin, nucleoli, nuclear membrane, endoplasmic reticulum, Golgi apparatus, vesicles, mitochondria, cytoplasm, cytoskeleton and membrane. We chose genes whose validated tagging terminus had at least one potential spacer nicking SpCas9’s non-target strand in each of four 15 bins within +/- 30 bp of the insertion site (start codon for N terminus or stop codon for C terminus). Hence, the prime editor could nick in the following bins: (-30 bp, -15 bp), (-15 bp, 0 bp), (0 bp, 15 bp), (15 bp, 30 bp). When multiple spacers were available in a bin we prioritized the spacer with the higher specificity determined by the number of off-target matches in the human genome through CRISPick, an sgRNA picker hosted by the Broad Institute. We prioritized specificity to minimize off-target tagging. Importantly, we did not consider predicted on-target scores of spacers a priori to be able to test more impartially the influence of on-target spacer activity on pegRNA activity. Lastly, we avoided spacers that had a polyT nucleotide stretch of > 4 bp due to longer polyT stretches known to drive significant termination of transcription by Pol III promoters, which could potentially cause dropout of pegRNAs due to poor pegRNA expression. Considering these design choices, we had four targeting spacers per gene for 60 genes total resulting in 240 targeting spacers total in the library.

We then used PrimeDesign^55^ to generate “parent” pegRNAs that we fed to a custom pipeline to generate design combinations that explored the following pegRNA parameters. For the PBS, we chose to test 10 nt, 13 nt and 16 nt lengths and for the RTT homology distal to the nick site we chose to test 15 nt, 27 nt, 36 nt and 48 nt. For the tag insertion, we tested three lengths 48 nt, 60 nt, 78 nt to test the generality of the method for different useful peptide lengths. Glycine-serine (GS) rich linkers of 4 aa and 10 aa lengths were coded within the RTT to be in-frame between the terminus and mNG(11) tag insertion for the 60 nt and 78 nt lengths respectively. In addition, for each pegRNA we incorporated a twin which was the same pegRNA except the tag insertion template had a synonymous codon sequence with maximal Hamming distance while avoiding long polyT stretches. Therefore, each pegRNA had an internal replicate that differed slightly in tag sequence and otherwise had the same composition for the other pegRNA components. To minimize re-nicking of the target DNA strand we mutated the PAM sequence (NGG) with silent mutations when possible or mutated NGG to NCC when silent mutations were not considered by PrimeDesign within the 5’ UTR or 3’ UTR of the targeted terminus. The libraries were split by insertion length into three sublibraries with 6000 pegRNAs including 240 non-targeting controls in each sublibrary.

The set of non-targeting controls was designed from 10 “parent” spacers that had no significant match to the human genome taken from the non-targeting controls hosted in the Broad Institute’s Genetic Perturbation Platform web portal. We generated a random target sequence with a protospacer for each “parent” non-targeting spacer and fed the target sequences to PrimeDesign to generate “parent” pegRNAs, which we subsequently fed to a custom pipeline to generate 24 pegRNA combinations for each “parent” spacer based on the pegRNA parameters noted earlier, resulting in 240 non-targeting pegRNA controls for each sublibrary. Importantly, these non-targeting pegRNA controls are not expected to lead to productive tagging events and should deplete from the library during sorting of tagged cell libraries.

### pegRNA pooled library cloning

To generate pooled pegRNA libraries compatible with single cell and optical sequencing approaches, we first modified the CROPSeq vector such that the full pegRNA and downstream barcode would be inserted between two BsmBI sites and not just the spacer sequence as in the original vector. Thus after the U6 promoter we had the first BsmBI site followed by a filler sequence, the second BsmBI site and the 3’ part of the padlock recognition sequence (ACTGGCTATTCATTCGC) followed by an RT primer binding site (CCTTTGGGTAAGCACACGTC) (Fig. 2B, plasmid deposited on Addgene). We then synthesized three pegRNA sublibraries corresponding to 48, 60 and 78nt tag lengths (see example sequences in Fig. S11) through Twist Biosciences. The oligo pool was amplified by PCR with custom sublibary-specific primers for the first two sublibraries of 48 and 60nt insertion lengths and hU6 forward primer and 3’ padlock reverse primer for the sublibrary with 78nt insertion (Fig. S11) following Twist oligo pool amplification manufacture instructions with 14 to 16 PCR cycles. In the first large scale cloning step, amplified oligo pools for 48 bp and 60 bp sublibraries were cloned using golden gate assembly with the BsmBI type II restriction enzyme and the 78 bp sublibrary was cloned using NEBuilder® HiFi DNA assembly. All three sublibraries were transformed using electrocompetent cells (Endura electrocompetent cells LGC) maintaining at least ∼300x coverage of transformed cells. Next, to insert the pegRNA scaffold, we performed a second large scale cloning step. A dsDNA fragment was produced by PCR (gttttagagctagaaatagcaagttaaaataaggctagtccgttatcaacttgaaaaagtggcaccgagtcggtgc) and cloned using a golden gate reaction using BbsI type II restriction enzyme for the 48 bp and 60 bp sublibraries and a BsmBI restriction enzyme for the 78 bp sublibrary. Electrocompetent cells (Endura electrocompetent cells LGC) were transformed with each library maintaining at least ∼100x coverage.

### Generation of pooled tagged libraries

To generate a pooled tagged cell library we first determined the titre of the lentiviral pegRNA library. This was done by transducing 2M HEK293T cells in each well of a 12 well plate using increasing volume of lentiviral supernatant. Cells were spinfected at suspension for 1 hour at 1000g at 32 degree celsius with 10 µg/mL polybrene (Millipore Sigma TR-1003). Cells are then passaged a day after spinfection to two new 12 well plates with and without puromycin selection (1 µg/mL) and selected for 4 days. Functional titre is determined as the ratio between the number of cells in the selected vs non selected population for each viral amount. To generate the cell libraries, multiple 12 well plates were transduced at a target MOI of less than 0.1 with the aim of achieving a coverage of 1000x of transduced cells for each sublibrary. Following transduction, cells were selected with puromycin (1 µg/mL) for seven days and continued to be expanded till day 23 from transduction. Tagged cells are then sorted using non tagged cells as a control for FACS gating (see Figure 2D). Sorting was performed at approximately 300x coverage for each sublibrary. Following expansion of sorted tagged cell libraries, a second round of sorting was performed to further purify the population of fluorescent cells.

### Bulk amplicon sequencing of pooled cell libraries

For analysis of pegRNA representation within the sorted and unsorted cell populations, cell pellets were collected with the number of cells exceeding 1000x coverage for each library. Genomic DNA (gDNA) was extracted using QIAamp DNA Mini Kit (QIAGEN Cat 51304) following manufacturer’s instructions. gDNA was quantified using Qubit broad range dsDNA kit (Thermo Cat Q32850). To amplify the pegRNA cassette with the downstream barcode from the integrated lentiviral cassettes, all extracted gDNA was used in 100ul PCR reactions of NEBNext Ultra II Q5 Master Mix (NEB Cat M0544X) using 10ug of gDNA as template and 22 cycles of exponential amplification. For forward primers a mix of staggered primers were used containing Illumina adapters and variable length sequence between the primer binding site and adapter. AATGATACGGCGACCACCGAGATCTACACTCTTTCCCTACACGACGCTCTTCCGATCT(stagger)TTGTGGAAAGGACGAAACACCG. For each sample a single barcoded reverse primer was used CAAGCAGAAGACGGCATACGAGAT (8bp barcode) GTGACTGGAGTTCAGACGTGTGCTCTTCCGATCTCAAAGGGCGAATGAATAGCCAGT. Resulting PCR products were mixed for each sample and ran on a 2% agarose gel. The appropriate size band was cut out of the gel and cleaned using the Monarch DNA Gel Extraction Kit Protocol (NEB Cat #T1020). For deep sequencing, barcoded samples were mixed at equal amounts and sequenced on a NextSeq2000 Illumina machine following manufacturer’s instructions.

### Analysis of epigenetic features

Epigenetic markers profiled for HEK293T and HEK293 (when HEK293T data was not available) were curated from publicly available datasets^23–33^. Corresponding bigwig files were accessed using pyBigWig and the average signal calculated across a 2 kb window centered on the start codon or stop codon insertion points of N terminus and C terminus targeted genes respectively. Averages of each mark were compared against average tagging efficiency for each gene.

### Modeling pegRNA tagging efficiencies

For each sequenced sample, pegRNA read counts were normalized for sequencing depth at reads per million, a pseudocount of 1 was added and counts were log2 transformed. Log2 fold changes were calculated for double sorted sublibraries relative to presort day 23 pegRNA normalized counts. For each sublibrary, pegRNAs were dropped if their normalized counts were at least two standard deviations below the sublibrary mean (z-score ≤ -2), resulting in 17,014 pegRNAs (16,295 targeting pegRNAs and 719 non-targeting pegRNA controls). For feature analysis, log2 fold changes of targeting pegRNAs were z-scored with respect to their cognate sublibrary distribution. For model training, z-scored log2 fold changes for each targeting pegRNA were mean-centered with respect to their cognate gene distribution across all sublibraries and pegRNAs were included if their targeted gene had at least one spacer with significant q-values (FDR < 0.1) for each sublibrary, resulting in the inclusion of 9,873 pegRNAs targeting 37 genes intersecting across all sublibraries.

XGBoost regression models^36^ were trained for prediction of pegRNA tagging efficiencies using nested cross-validation for hyperparameter tuning and model evaluation. A leave-one-gene-out (LOGO) strategy was used for evaluating outer folds and grouped k-fold cross-validation (k=6) was used for hyperparameter tuning within inner folds. The optimal hyperparameter combination for each outer fold was found through grid search and defined as the combination with the minimum average mean squared error (MSE) regression loss across inner folds. Hyperparameters tuned were: number of trees (50, 100, 150), max depth (3, 4, 5), learning rate (0.01, 0.05, 0.1), minimum loss reduction (0, 0.01, 0.1), L1 regularization (0, 0.01, 0.1) and L2 regularization (0.01, 0.1, 1). Model evaluation metrics were: average spearman correlation between predicted and observed tagging efficiencies and average precision at top k for a given gene within a sublibrary. A model for deployment was subsequently trained using the most frequent, optimal hyperparameter combination found across outer folds. Feature contributions to model predictions were assessed through SHAP (SHapley Additive exPlanations) value analysis^38^.

For initial model training and evaluation, 63 pegRNA features were used as inputs, including: on-target spacer efficacy score (Rule Set 3)^35^, nicked strand, nick position relative to insertion point, nick distance from insertion point, PAM disruption, first nucleotide of the 3’ extension as well as length, nucleotide content, GC content, melting temperature and mean free energy^37^ for different regions of the pegRNA. Iterative feature selection was performed by iteratively building models, selecting input features with non-zero mean absolute SHAP values and feeding them as inputs to the next model iteration until only features with non-zero mean absolute SHAP values remained. The final model for deployment used the consolidated set of 37 features and trained with the following hyperparameters: number of trees = 50, max depth = 3, learning rate = 0.05, minimum loss reduction = 0, L1 regularization = 0.1 and L2 regularization = 1. Each model iteration was tuned, evaluated and analyzed as noted earlier.

Python packages: native XGBoost Learning API, xgboost, was used for model training, joblib was used to parallelize hyperparameter tuning across multiple cores, sklearn to calculate mean square error, scipy to calculate spearman correlations and SHAP for model feature analysis.

### Cell imaging and in-situ sequencing of pegRNA spacers and barcodes

A glass 6-well plate (CellVis, P06-1.5H-N) was coated with fibronectin (Corning 354008) at 50 ug/ml for two hours at RT and pooled tagged cell libraries were plated at a seeding density of 700,000 cells per well. 18 to 24 hours post-plating the wells were stained with Syto85 at 100 nM (ThermoFisher S11366) and incubated for 30 minutes at 37°C with 5% CO2. Next, wells were fixed with 4% PFA (Electron Microscopy Services 15710) for 30 minutes at RT followed by actin and nuclei staining with Alexa Fluor Plus 750 Phalloidin diluted to 0.1X (ThermoFisher A30105) and DAPI at 266 ng/ml (ThermoFisher 62248) respectively in an RNAse-free PBS solution (ThermoFisher AM9625) containing Ribolock RNAse inhibitor at 0.8 U/ul (ThermoFisher, EO0382) for 1 hour at RT. The stained plate was phenotyped with an inverted Ti2 Nikon Eclipse microscope outfitted with an X-Light V3 (CrestOptics) for control of filters and dichroic mirrors and a CELESTA 7 channel light engine (Lumencor, 90-10525) for sample illumination. Phenotyping images were collected on an ORCA-Fusion Gen-III sCMOS camera (Hamamatsu C14440-20UP) with channel-specific filters at 40X magnification using the following settings: DAPI: 408 nm laser, 10% power, 20 ms exposure time; mNeonGreen: 477 nm laser, 30% power, at two exposure times 100 ms and 300 ms, Syto85: 546 nm laser, 5% power, 80 ms exposure time, AlexaFluor750-Phalloidin: 749 nm laser, 30% power, 200 ms exposure time. An automated JOB programmed in Nikon Elements (v5.21.03) was used to tile each well and capture images of size 2304 x 2304 pixels at 40X magnification (0.17 mpp, total of 35,118 images for a 6-well plate). DAPI nuclear stain was used for nuclear segmentation and Phalloidin actin stain for cellular segmentation.

After phenotyping, we proceeded with a standard in-situ sequencing workflow^41^ with the following exception: we added a pre-RT fixation step with amine-modified RT primers. We performed 12 cycles of in-situ sequencing at 10X magnification to read out the pegRNA spacer for three wells (each of these wells corresponded to a distinct sublibrary) and the 12 bp barcode representing the entire pegRNA for the other three wells (all of these wells corresponded to the 60 bp insertion sublibrary) using CROPseq and LentiGuide-BC probe sets^40^ respectively (sequences noted below). Each sequencing cycle was performed by using an automated Nikon Elements JOB to tile each well and capture images of size 2304 x 2304 pixels at 10X magnification (0.64 mpp, total of 2,172 images for a 6-well plate). The following settings were used for genotyping: DAPI: 408 nm laser, 5% power, 50 ms exposure time; G and T: 546 nm laser, 30% power, 200 ms exposure time; A: 546 nm laser, 30% power, 200 ms exposure time; and C: 546 nm laser, 30% power, 200 ms exposure time. Images contained 5 channels per sequencing cycle: DAPI nuclear stain for alignment between cycles and nucleotide imaging channels G, T, A and C for sequencing.

### CROPseq probe set to in-situ sequence pegRNA spacers

RT primer: /5AmMC12/G+AC+TA+GC+CT+TA+TT+TTAACTTGCTAT

Padlock:

/5Phos/gttttagagctagaaatagcaagCTCCTGTTCGACACCTACCCACCTCATCCCACTCTTCAaaag gacgaaacaccg

Sequencing primer: CACCTCATCCCACTCTTCAaaaggacgaaacaccg

### LentiGuide-BC probe set to in-situ sequence pegRNA barcodes RT primer: /5AmMC12/G+AC+GT+GT+GC+TT+AC+CCAAAGG

Padlock:

/5Phos/actggctattcattcgcCTCCTGTTCGACAGTCAGCCGCATCTGCGTCTATTTAGTGGAGCCCTTGtgttcaatcaacattcc

Sequencing primer: TTCGACAGTCAGCCGCATCTGCGTCTATTTAGTGGAGCCCTTGtgttcaatcaacattcc

The genotyping pipeline published by Feldman, Singh et al. (2019)^41^ was adapted for genotyping analysis. Images across all cycles were assembled for every tile to produce a 4 dimensional array (cy x ch x h x w), every tile was analyzed independently. Sequencing peaks are identified, and attributed a nucleotide read for every cycle based on the brightness of sequencing channels. A specific spacer from the library is assigned to a cell based on the barcodes obtained from sequencing peaks. Spacers are assigned to cells with at least 3 peaks that agree on the same barcode. Every sequencing peak is assigned to an individual cell using the segmented cellular masks. Nuclei and cells were segmented using Cellpose^56^. Nuclei were segmented using the DAPI channel, using single channel segmentation with Cellpose ‘nuclei’ model. Cells were segmented using two channel segmentation, DAPI for nuclei and C channel for cellular, with Cellpose ‘cyto2’ model. Cell and nuclei masks were then consolidated such that both nuclear and cellular masks share the same label. Cells touching image boundaries, nuclei with no cell, and cells with no or multiple nuclei were eliminated from the tile’s cellular and nuclear masks.

### Image analysis and generation of cell albums

Single cell phenotyping data was calculated for every cell including, nuclear area, minimum, maximum, mean, and standard deviation of intensity of all imaging channels. Phenotyping images of cells were linked to genotyping images using a mapping function. Nuclear mask centroids at 10X images were mapped to nuclear mask centroids at 40X using an affine transformation with 5 degrees of freedom including x-translation, y-translation, angle of rotation, x-scaling, and y-scaling,

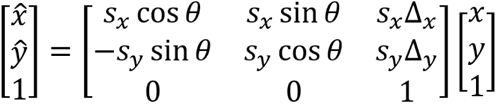

The optimal degrees of freedom were calculated using 5 fiducial cells obtained manually, and used as targets for a minimization algorithm,

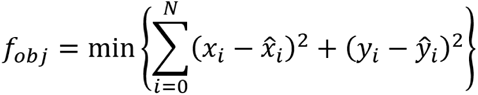

Single cell images with masks were produced for further analysis once cells’ phenotype and genotype were linked. Cells tagged with the same spacer were combined into a cell picture album to depict the spacer’s protein localization pattern for quality control and manual inspection.

### Training of cytoself VAE and analysis of latent parameters

Cytoself^45^ (pytorch version) variational autoencoder (VAE) was trained on single cell images from 103 sgRNAs tagging 50 genes. The average intensity of single cell images were binned into a log-histogram for each sgRNA . A gaussian filter was used to smooth the histogram and identify the mean of the histogram distribution. Cells were selected from within one standard deviation of the mean. In the case of multiple peaks, the peak with higher intensity was used. A maximum of 1500 cells were selected for each of 103 sgRNAs. Cytoself dataset includes three channels, first channel contains the entire cell’s phenotype cropped using the cellular mask, second channel contains only the nuclear portion of the cells cropped using the nuclear mask, third channel contains only the cytosolic portion of the cell, cropped by subtracting the nuclear mask from the cellular mask. Raw cell images were resized to 128 x 128 shape and assembled into a single array. Training was done on a Nvidia A100-80GB GPU with a batch size of 256. Training, validation, test sets were split 0.7, 0.15, and 0.15 respectively. The global representation latent space was reduced to a 2D UMAP, and clustered by the unsupervised clustering algorithm HDBSCAN, using a minimum cluster size of 600 and epsilon of 0.7. Training parameters were used as follows (Input Shape - 3 x 128 x 128, Embedding Shape - (32, 32), (4, 4), Embedding Dimensions - 64, Embedding Number - 512, FC Output Index - 2, Class Number - 12, FC Input Type - vqindhist, Learning rate - 1e-4, Max Epoch - 100, Reduce Learning Rate Patience - 3, Reduce Learning Rate Increment - 0.1, Early Stop Patience - 10).

## Bibliography

1. Cho, N. H. et al. OpenCell: Endogenous tagging for the cartography of human cellular organization. Science 375, eabi6983 (2022).

2. Schmid-Burgk, J. L., Höning, K., Ebert, T. S. & Hornung, V. CRISPaint allows modular base-specific gene tagging using a ligase-4-dependent mechanism. Nat. Commun. 7, 12338 (2016).

3. Kim, J., et al. High-throughput tagging of endogenous loci for rapid characterization of protein function bioRxiv 2022.11.16.516691 (2023) doi:10.1101/2022.11.16.516691.

4. Serebrenik, Y. V., Sansbury, S. E., Kumar, S. S., Henao-Mejia, J. & Shalem, O. Efficient and flexible tagging of endogenous genes by homology-independent intron targeting. Genome Res. 29, 1322–1328 (2019).

5. Reicher, A., Koren, A. & Kubicek, S. Pooled protein tagging, cellular imaging, and in situ sequencing for monitoring drug action in real time. Genome Res. 30, 1846–1855 (2020).

6. Sansbury, S. E., Serebrenik, Y. V., Lapidot, T., Burslem, G. M. & Shalem, O. Pooled tagging and hydrophobic targeting of endogenous proteins for unbiased mapping of unfolded protein responses. bioRxiv (2023) doi:10.1101/2023.07.13.548611.

7. Serebrenik, Y. V., Mani, D., Maujean, T., Burslem, G. M. & Shalem, O. Pooled endogenous protein tagging and recruitment for scalable discovery of effectors for induced proximity therapeutics. bioRxiv (2023) doi:10.1101/2023.07.13.548759.

8. Anzalone, A. V. et al. Search-and-replace genome editing without double-strand breaks or donor DNA. Nature 576, 149–157 (2019).

9. Chen, P. J. & Liu, D. R. Prime editing for precise and highly versatile genome manipulation. Nat. Rev. Genet. 24, 161–177 (2023).

10. Kim, H. K. et al. Predicting the efficiency of prime editing guide RNAs in human cells. Nat. Biotechnol. 39, 198–206 (2021).

11. Yu, G. et al. Prediction of efficiencies for diverse prime editing systems in multiple cell types. Cell 186, 2256–2272.e23 (2023).

12. Mathis, N. et al. Predicting prime editing efficiency and product purity by deep learning. Nat. Biotechnol. 41, 1151–1159 (2023).

13. Koeppel, J. et al. Prediction of prime editing insertion efficiencies using sequence features and DNA repair determinants. Nat. Biotechnol. 41, 1446–1456 (2023).

14. Ren, X. et al. High-throughput PRIME-editing screens identify functional DNA variants in the human genome. Mol. Cell 83, 4633–4645.e9 (2023).

15. Erwood, S. et al. Saturation variant interpretation using CRISPR prime editing. Nat. Biotechnol. 40, 885–895 (2022).

16. Leonetti, M. D., Sekine, S., Kamiyama, D., Weissman, J. S. & Huang, B. A scalable strategy for high-throughput GFP tagging of endogenous human proteins. Proc. Natl. Acad. Sci. U. S. A. 113, E3501–8 (2016).

17. Kamiyama, D. et al. Versatile protein tagging in cells with split fluorescent protein. Nat. Commun. 7, 11046 (2016).

18. Zhou, S., Feng, S., Brown, D. & Huang, B. Improved yellow-green split fluorescent proteins for protein labeling and signal amplification. PLoS One 15, e0242592 (2020).

19. Matlashov, M. E. et al. A set of monomeric near-infrared fluorescent proteins for multicolor imaging across scales. Nat. Commun. 11, 239 (2020).

20. Doench, J. G. et al. Optimized sgRNA design to maximize activity and minimize off-target effects of CRISPR-Cas9. Nat. Biotechnol. 34, 184–191 (2016).

21. Hanna, R. E. & Doench, J. G. A case of mistaken identity. Nat. Biotechnol. 36, 802–804 (2018).

22. Hegde, M., Strand, C., Hanna, R. E. & Doench, J. G. Uncoupling of sgRNAs from their associated barcodes during PCR amplification of combinatorial CRISPR screens. PLoS One 13, e0197547 (2018).

23. Tak, Y. E. et al. Augmenting and directing long-range CRISPR-mediated activation in human cells. Nat. Methods 18, 1075–1081 (2021).

24. ENCODE Project Consortium. An integrated encyclopedia of DNA elements in the human genome. Nature 489, 57–74 (2012).

25. Luo, Y. et al. New developments on the Encyclopedia of DNA Elements (ENCODE) data portal. Nucleic Acids Res. 48, D882–D889 (2020).

26. Hitz, B. C., et al. The ENCODE Uniform Analysis Pipelines. bioRxiv (2023) doi:10.1101/2023.04.04.535623.

27. Thurman, R. E. et al. The accessible chromatin landscape of the human genome. Nature 489, 75–82 (2012).

28. Natarajan, A., Yardimci, G. G., Sheffield, N. C., Crawford, G. E. & Ohler, U. Predicting cell-type-specific gene expression from regions of open chromatin. Genome Res. 22, 1711–1722 (2012).

29. Wu, T., Lyu, R., You, Q. & He, C. Kethoxal-assisted single-stranded DNA sequencing captures global transcription dynamics and enhancer activity in situ. Nat. Methods 17, 515–523 (2020).

30. Liang, K. et al. Targeting Processive Transcription Elongation via SEC Disruption for MYC-Induced Cancer Therapy. Cell 175, 766–779.e17 (2018).

31. Schachner, L. F. et al. Decoding the protein composition of whole nucleosomes with Nuc-MS. Nat. Methods 18, 303–308 (2021).

32. Morgan, M. A. J. et al. A cryptic Tudor domain links BRWD2/PHIP to COMPASS-mediated histone H3K4 methylation. Genes Dev. 31, 2003–2014 (2017).

33. Fan, H. et al. BAHCC1 binds H3K27me3 via a conserved BAH module to mediate gene silencing and oncogenesis. Nat. Genet. 52, 1384–1396 (2020).

34. Li, X., et al. Chromatin context-dependent regulation and epigenetic manipulation of prime editing. bioRxiv (2023) doi:10.1101/2023.04.12.536587.

35. DeWeirdt, P. C. et al. Accounting for small variations in the tracrRNA sequence improves sgRNA activity predictions for CRISPR screening. Nat. Commun. 13, 5255 (2022).

36. Chen, T. & Guestrin, C. XGBoost: A Scalable Tree Boosting System. in Proceedings of the 22nd ACM SIGKDD International Conference on Knowledge Discovery and Data Mining 785–794 (Association for Computing Machinery, New York, NY, USA, 2016).

37. Lorenz, R. et al. ViennaRNA Package 2.0. Algorithms Mol. Biol. 6, 26 (2011).

38. Lundberg, S. & Lee, S.-I. A Unified Approach to Interpreting Model Predictions. arXiv [cs.AI] (2017).

39. Lundberg, S. M. et al. From Local Explanations to Global Understanding with Explainable AI for Trees. Nat Mach Intell 2, 56–67 (2020).

40. Datlinger, P. et al. Pooled CRISPR screening with single-cell transcriptome readout. Nat. Methods 14, 297–301 (2017).

41. Feldman, D. et al. Optical Pooled Screens in Human Cells. Cell 179, 787–799.e17 (2019).

42. Feldman, D. et al. Pooled genetic perturbation screens with image-based phenotypes. Nat. Protoc. 17, 476–512 (2022).

43. Labitigan, R. L. D. et al. Mapping variation in the morphological landscape of human cells with optical pooled CRISPRi screening. bioRxiv 2022.12.27.522042 (2022) doi:10.1101/2022.12.27.522042.

44. Bar-Even, A. et al. Noise in protein expression scales with natural protein abundance. Nat. Genet. 38, 636–643 (2006).

45. Kobayashi, H., Cheveralls, K. C., Leonetti, M. D. & Royer, L. A. Self-supervised deep learning encodes high-resolution features of protein subcellular localization. Nat. Methods 19, 995–1003 (2022).

46. Clarke, R. et al. Enhanced Bacterial Immunity and Mammalian Genome Editing via RNA-Polymerase-Mediated Dislodging of Cas9 from Double-Strand DNA Breaks. Mol. Cell 71, 42–55.e8 (2018).

47. Duan, M., Speer, R. M., Ulibarri, J., Liu, K. J. & Mao, P. Transcription-coupled nucleotide excision repair: New insights revealed by genomic approaches. DNA Repair 103, 103126 (2021).

48. Yarnall, M. T. N. et al. Drag-and-drop genome insertion of large sequences without double-strand DNA cleavage using CRISPR-directed integrases. Nat. Biotechnol. 41, 500–512 (2023).

49. Anzalone, A. V. et al. Programmable deletion, replacement, integration and inversion of large DNA sequences with twin prime editing. Nat. Biotechnol. 40, 731–740 (2022).

50. Wang, J. et al. Efficient targeted insertion of large DNA fragments without DNA donors. Nat. Methods 19, 331–340 (2022).

51. Zheng, C. et al. Template-jumping prime editing enables large insertion and exon rewriting in vivo. Nat. Commun. 14, 3369 (2023).

52. Wu, K. et al. De novo design of modular peptide-binding proteins by superhelical matching. Nature 616, 581–589 (2023).

53. Mercer, J. A. M. et al. Continuous evolution of compact protein degradation tags regulated by selective molecular glues. Science 383, eadk4422 (2024).

54. Kudo, T. et al. Highly multiplexed, image-based pooled screens in primary cells and tissues with PerturbView. bioRxiv 2023.12.26.573143 (2023) doi:10.1101/2023.12.26.573143.

55. Hsu, J. Y. et al. PrimeDesign software for rapid and simplified design of prime editing guide RNAs. Nat. Commun. 12, 1034 (2021).

56. Stringer, C., Wang, T., Michaelos, M. & Pachitariu, M. Cellpose: a generalist algorithm for cellular segmentation. Nat. Methods 18, 100–106 (2021).

