## Supplementary Fig 1-10 for "High-throughput optimized prime editing mediated endogenous protein tagging for pooled imaging of protein localization"

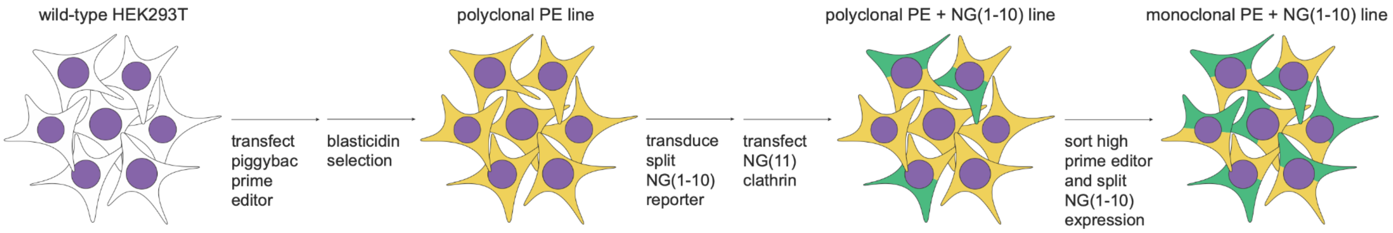


**Figure S1 - Illustration of how clonal lines for tagging were generated.** Wild-type cells were co-transfected with piggybac prime editor (PE2) and piggybac transposase followed by blasticidin selection. We then transduced the cells with a lentiviral vector containing split $mNG3K_{1-10}$ (addgene #157993) and transfected a plasmid expressing CLTA tagged with $mNG2_{11}$ to produce a fluorescence signal that would indicate the expression of the $mNG3K_{1-10}$ cassette. Cells where then sorted based on expression of both PE2 (far red channel) and $mNG3K_{1-10}$ (green channel).

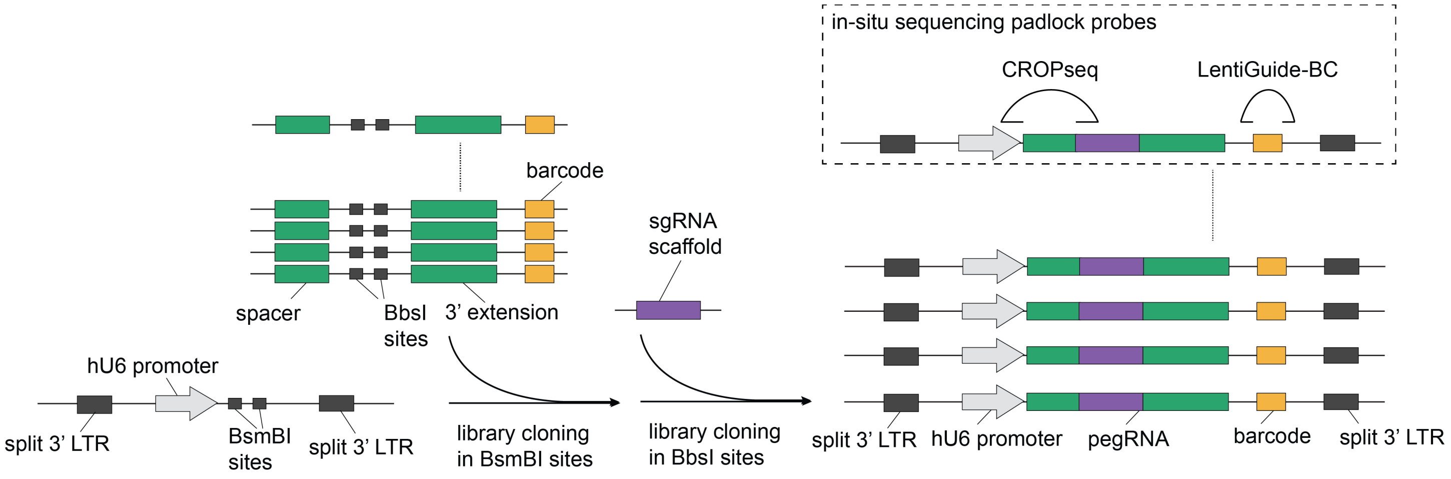


**Figure S2 - Illustration of the cloning strategy for pooled pegRNA libraries within a modified CROPseq vector.** See methods for a more detailed description of the cloning strategy. CROPseq vector was modified such that the original BsmBI restriction sites are not followed by a constant sgRNA scaffold but instead contain a 3' padlock hybridization sequence and RT primer site which are required for in-situ sequencing. Library cloning was performed by synthesizing an oligo pool containing pegRNAs with the constant sgRNA scaffold replaced by BbsI sites for 48nt and 60nt length sublibraries and BsmBI sites for the 78nt length sublibrary. The first cloning step was performed by golden gate for 48nt and 60nt length sublibraries and with Gibson cloning for the longer 78nt sublibrary. A second step of large scale golden gate cloning was performed to insert the constant sgRNA scaffold using corresponding BsmBI or BbsI sites as entry points. In-situ sequencing padlock probes are shown within inset: CROPseq padlock probe can be used to read out pegRNA spacer and LentiGuide-BC padlock probe can be used to read out the barcode.

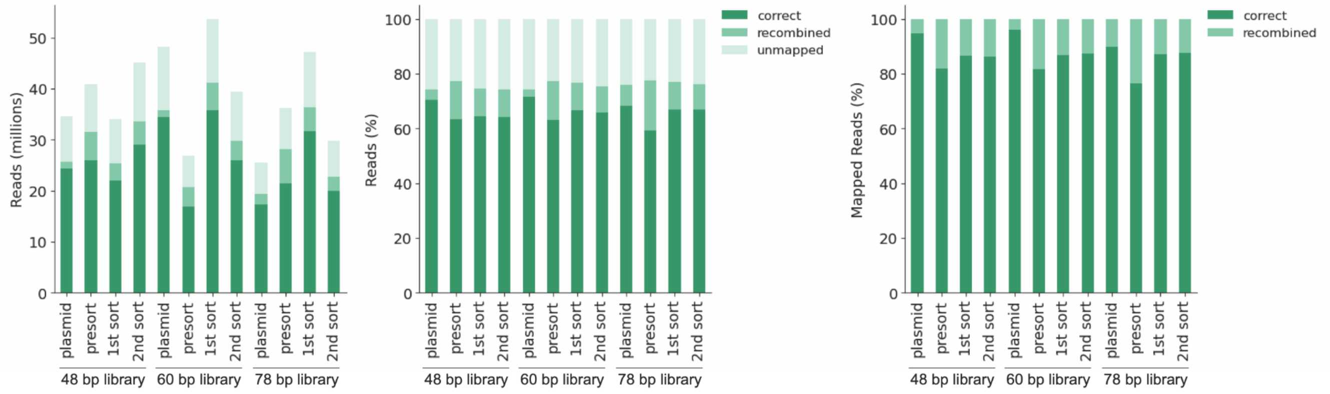


**Figure S3 - Read mapping and spacer-barcode recombination in different library stages.** Bar plots showing the number of reads (left panel), percent of reads (middle panel) and percent out of mapped reads (right panel) of correct and recombined sequences.

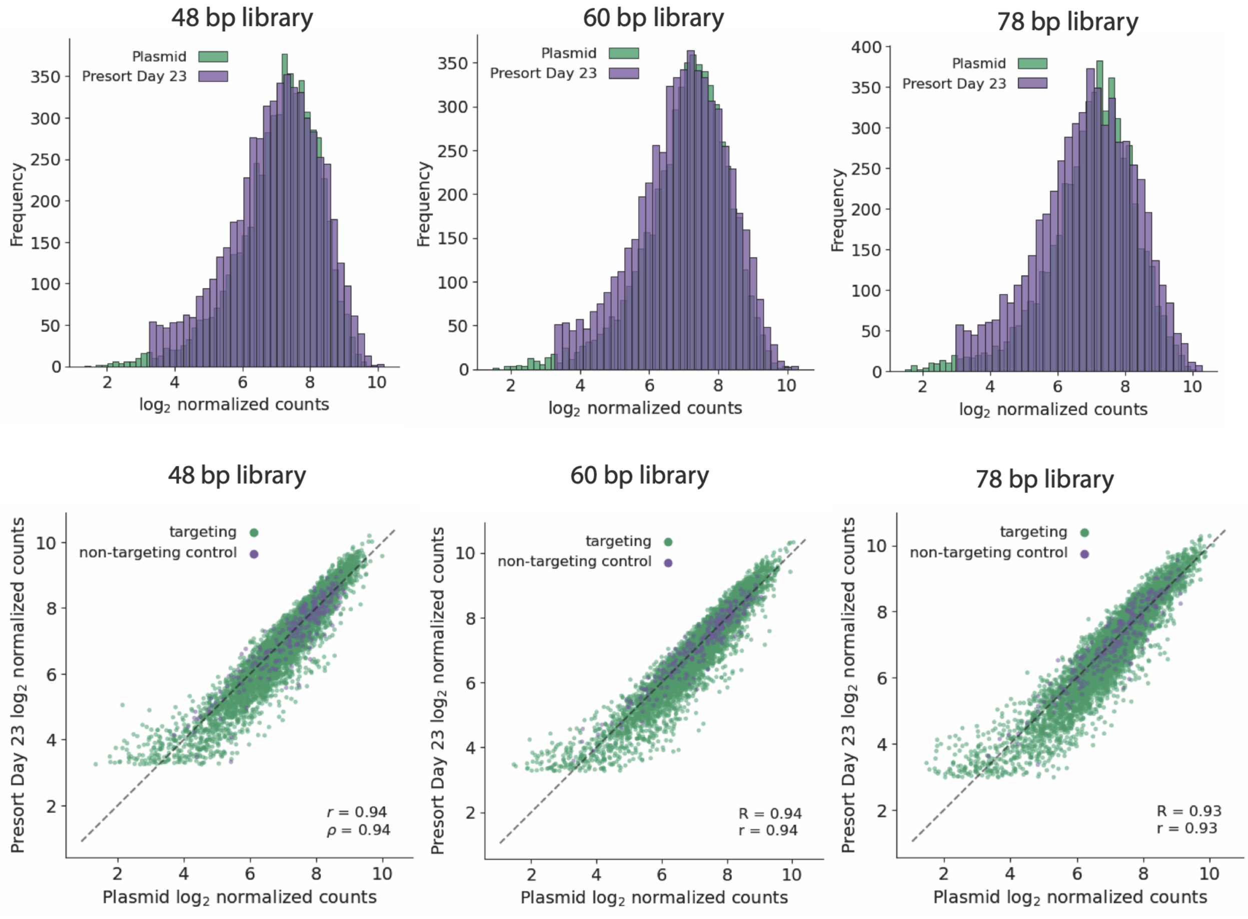


**Figure S4 - Read distributions for the three insertion length libraries after cloning and day 23 post-transduction before sorting.** Upper panels show histograms of read distributions demonstrating good coverage and log-normal distribution. Lower panels show comparisons between plasmid libraries and presort day 23 post-transduced cell libraries. Each scatterplot is annotated with Pearson r and Spearman R correlation coefficients.

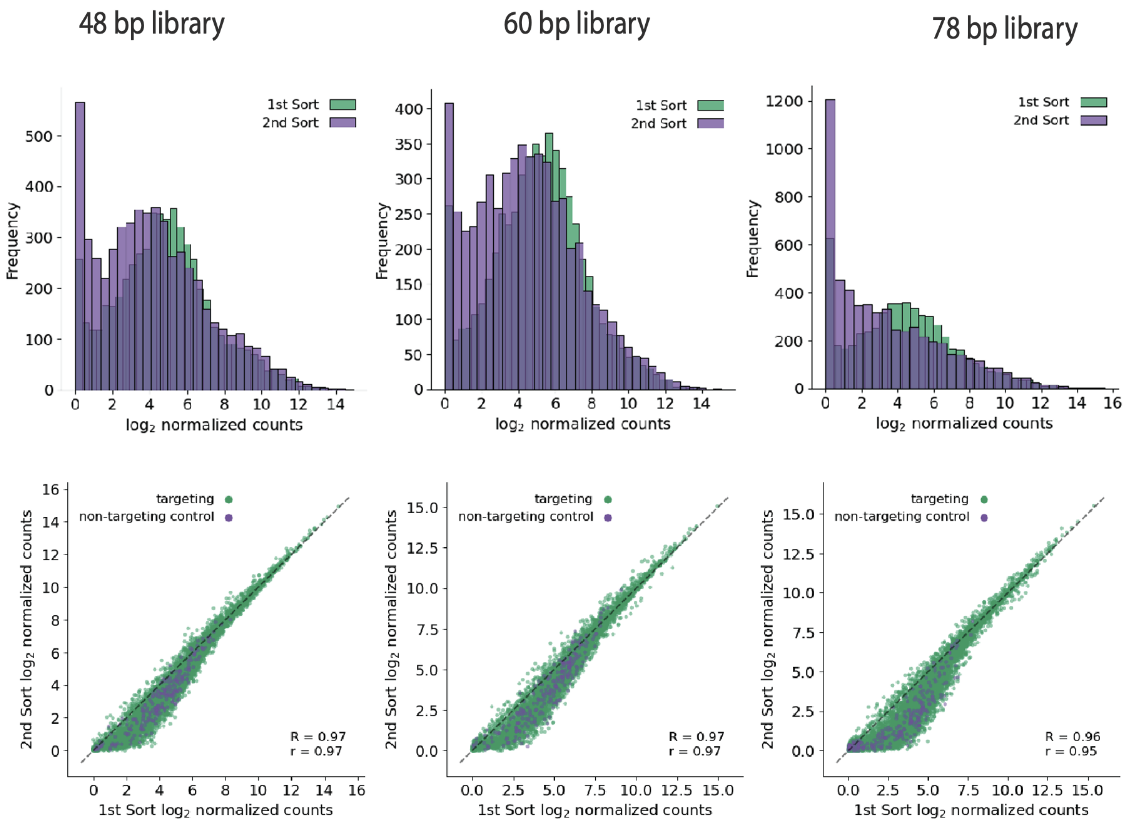


**Figure S5 - Read distributions for the three insertion length cell libraries after sorting fluorescent cells.** Upper panels show histograms of read distributions demonstrating depletion of pegRNAs that resulted in no gene tagging and increased library skew due to sorting and different tagging efficiencies between genes. Lower panels compare 1$^{st}$ round and 2$^{nd}$ round sorted libraries. Each scatterplot is annotated with Pearson r and Spearman R correlation coefficients.

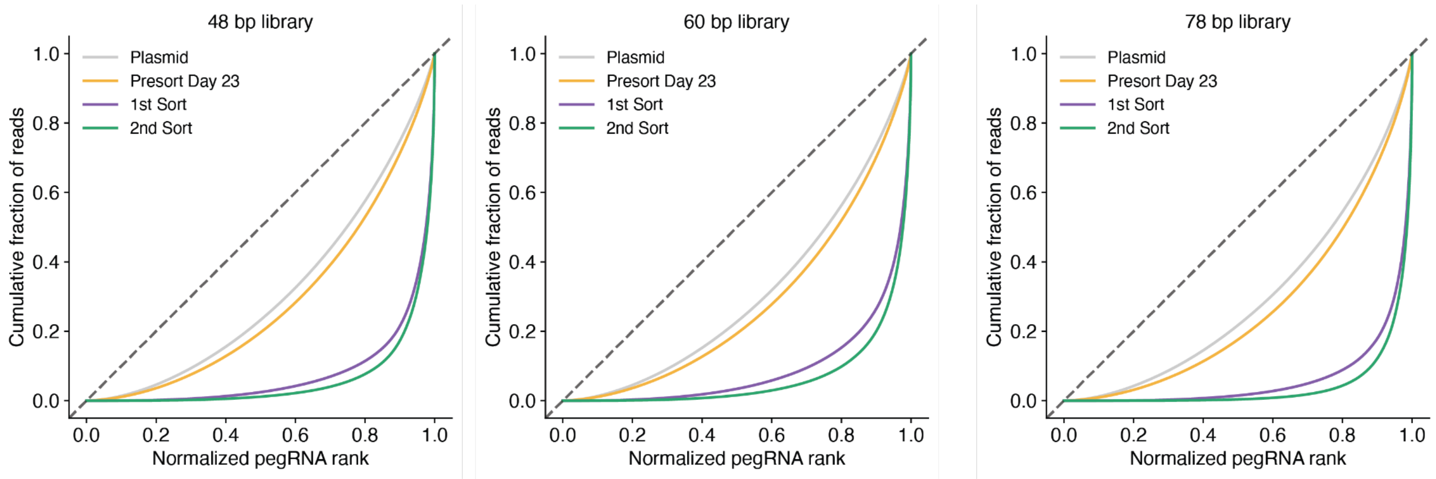


**Figure S6 - Lorenz curves for each library across samples showing increased skew from plasmid libraries to sorted libraries.** Increased library skew is particularly notable in sorted libraries due to depletion of pegRNAs that resulted in no gene tagging and differential tagging efficiency between genes.

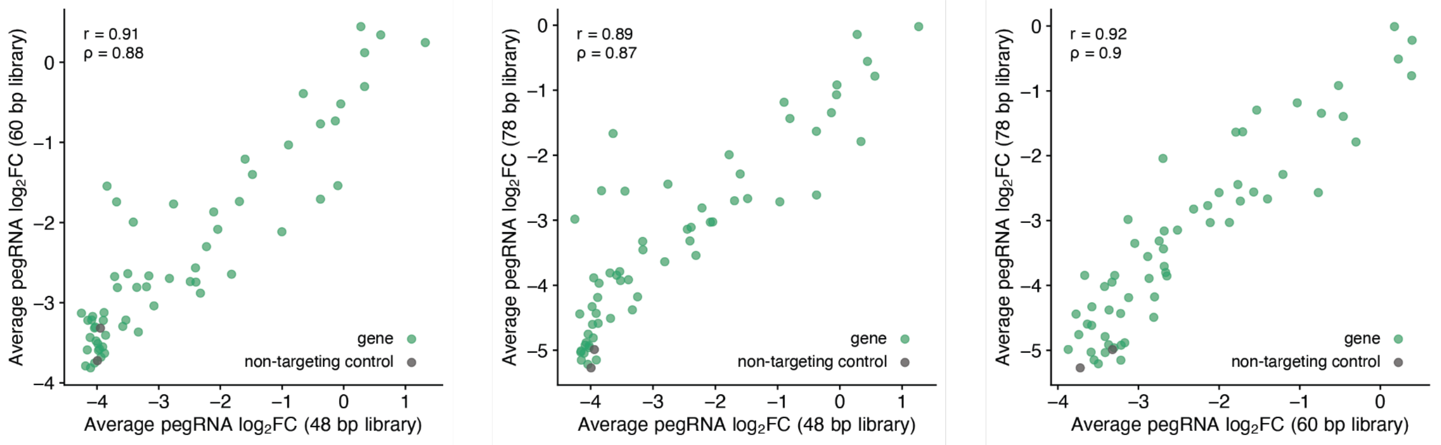


**Figure S7 - Correlations of average gene effect between libraries with different insertion sizes.** Pairwise comparison of average pegRNA $\log_2$FC for each gene across three insertion libraries. Green dots indicate genes targeted and black dots indicate non-targeting control pseudo-genes. Each scatterplot is annotated with Pearson r and Spearman ρ correlation coefficients.

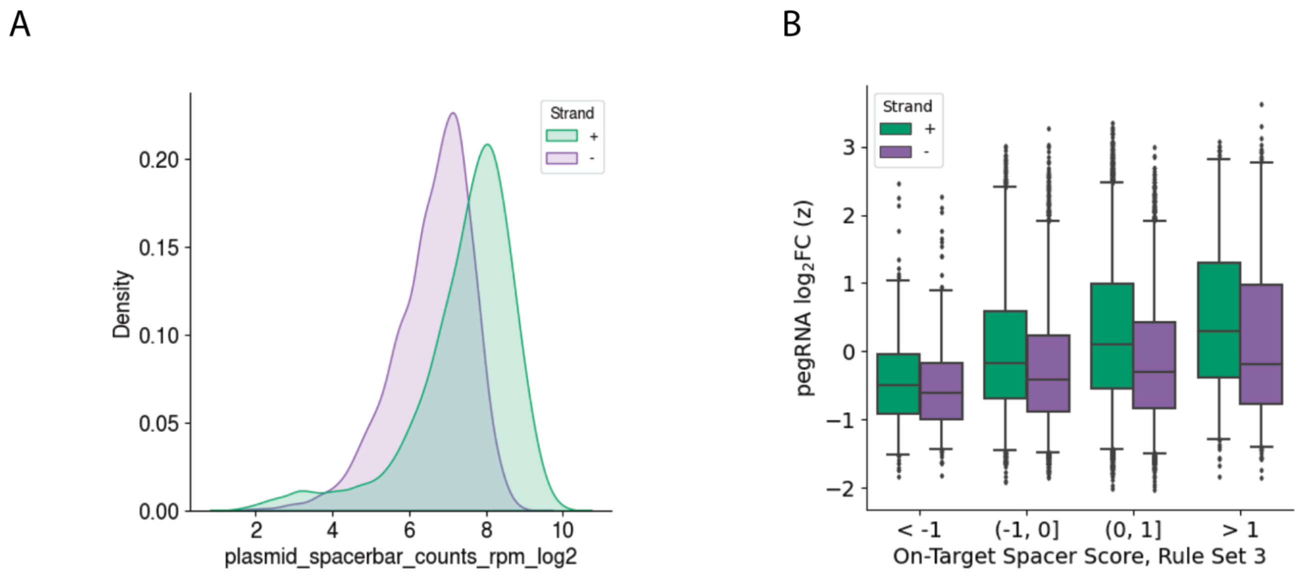


**Figure S8 - Nicked strand effects on pegRNA tagging efficiency and initial abundance.** (**A**) Comparison of the abundance distribution of +/- strand nicking pegRNAs in the initial cloned plasmid library. The difference is likely due to differential oligo synthesis or PCR amplification as +/- strand nicking pegRNAs encode two distinct insertion sequences with the peptide insertion template in either coding or reverse complement orientations. (**B**) pegRNAs are binned based on on-target spacer score (RS3) showing increased tagging efficiency at the + strand independent of spacer cutting score. In each strand legend, + indicates coding strand and - indicates template strand.

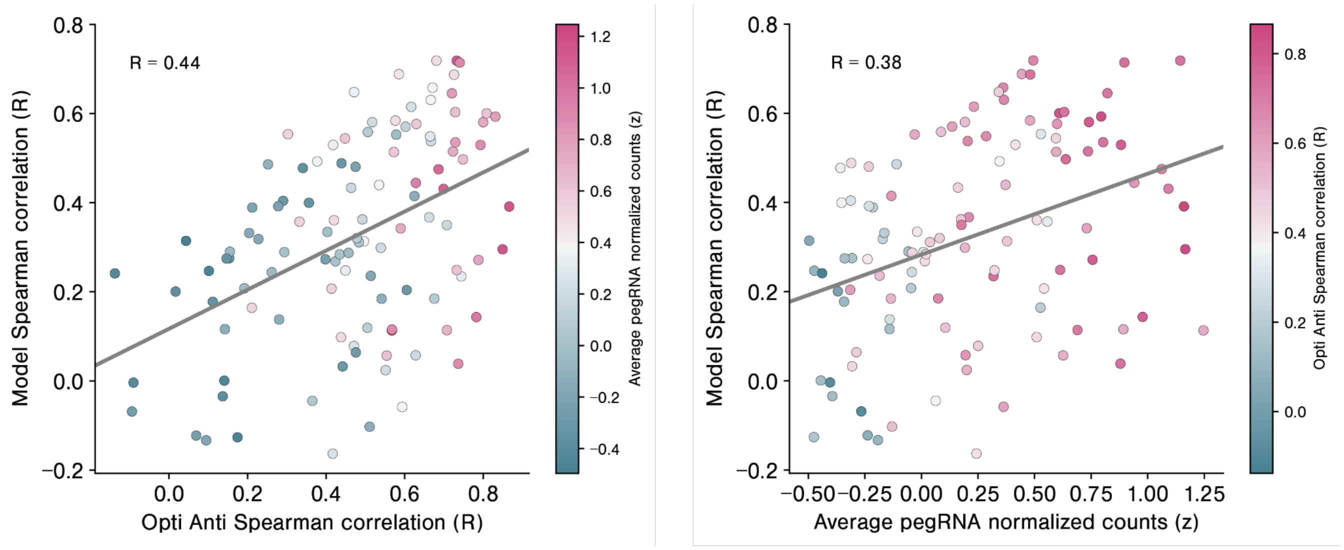


**Figure S9 - Comparison between model performance and pegRNA tagging reproducibility.** (**A**) Correlation between model performance (y axis) with pegRNA tagging reproducibility (x axis). (**B**) Correlation between model performance (y axis) and pegRNA abundance (x axis). Each scatterplot is annotated with Spearman R correlation coefficients. Each dot is a gene within a sublibrary.

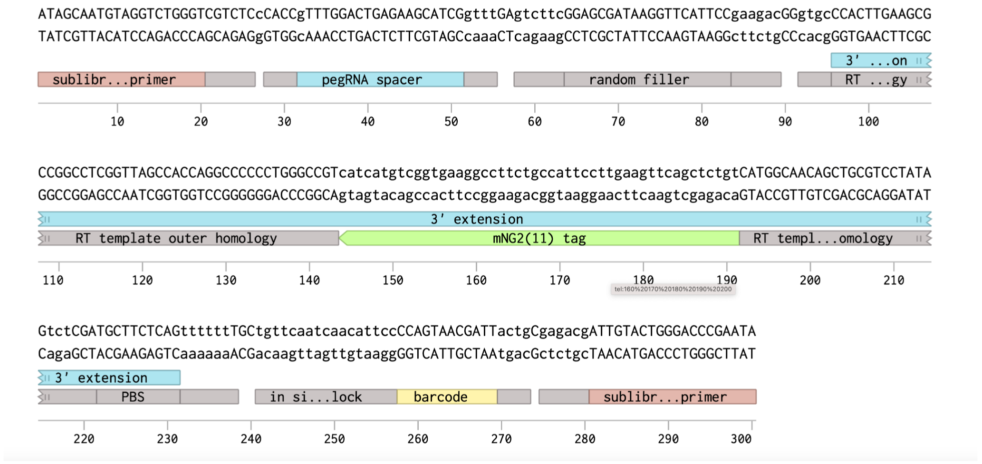


48 bp length

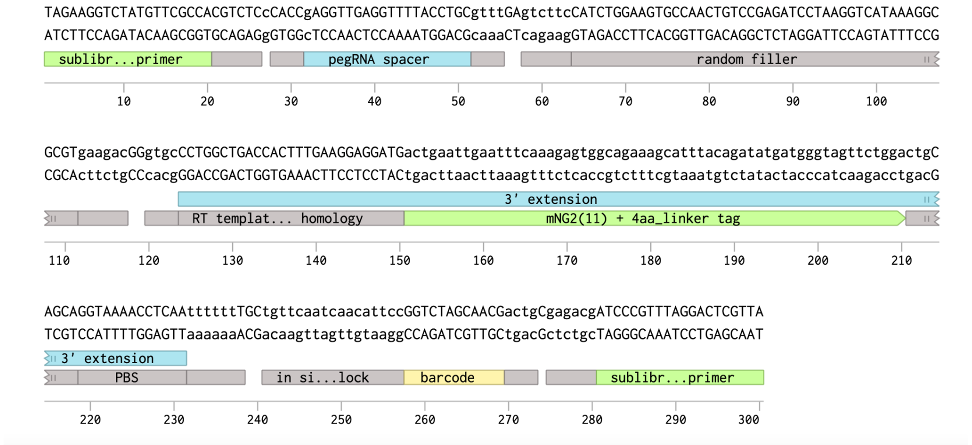


60 bp length

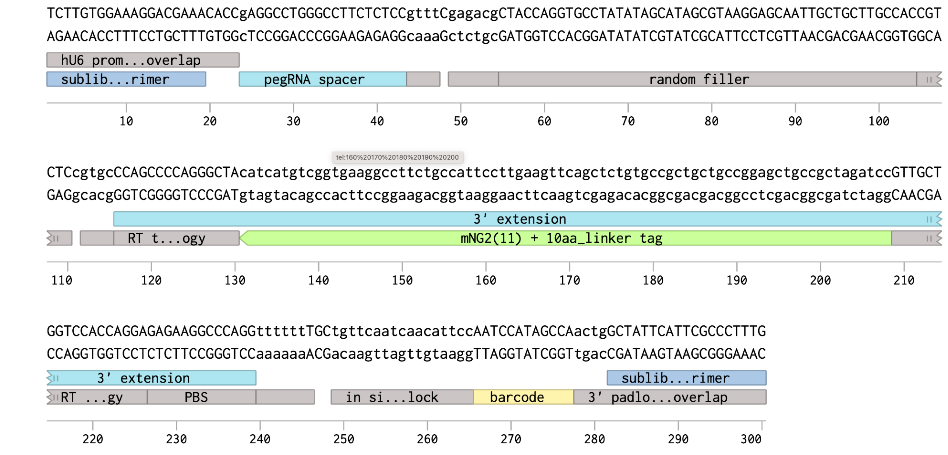


78 bp length

**Figure S10 - Examples of synthesized pegRNAs in the oligo pool library for the three insertion lengths.** Parts of the synthesized oligo from left to right (48 bp insertion as example): fw primer for sublibrary amplification (1-20), BsmBI site for golden gate cloning (21-26), pegRNA spacer with flanking golden gate overlaps (28-55), random filler flanked by BbsI sites (58-89), pegRNA 3' extension (92-231), U6 terminator (232-238), in situ 5' padlock (241-257), barcode (258-269), BsmBI site for golden gate cloning (275-280), rv primer for sublibrary amplification (281-300).
