## Supplementary figures and images for "High-throughput optimized prime editing mediated endogenous protein tagging for pooled imaging of protein localization"

### Figure S11

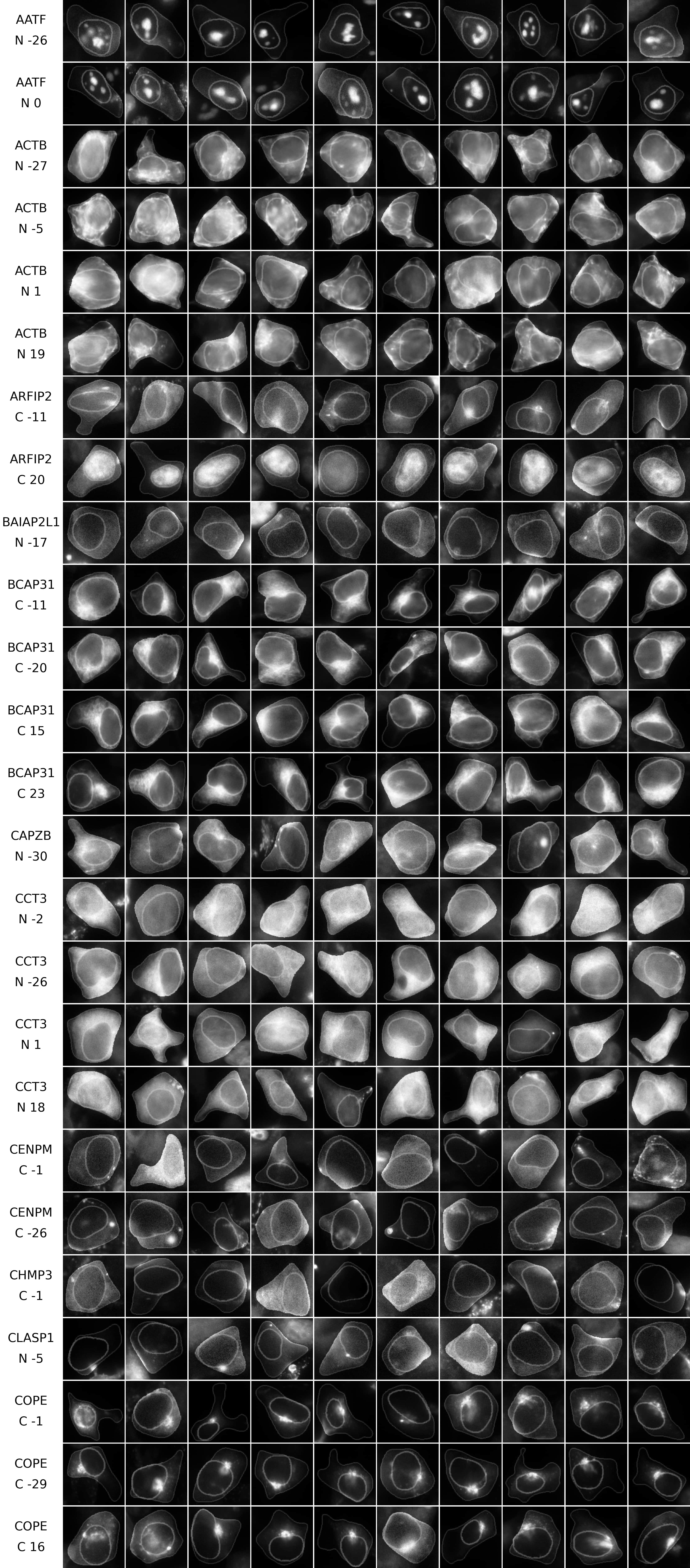

### Figure S12

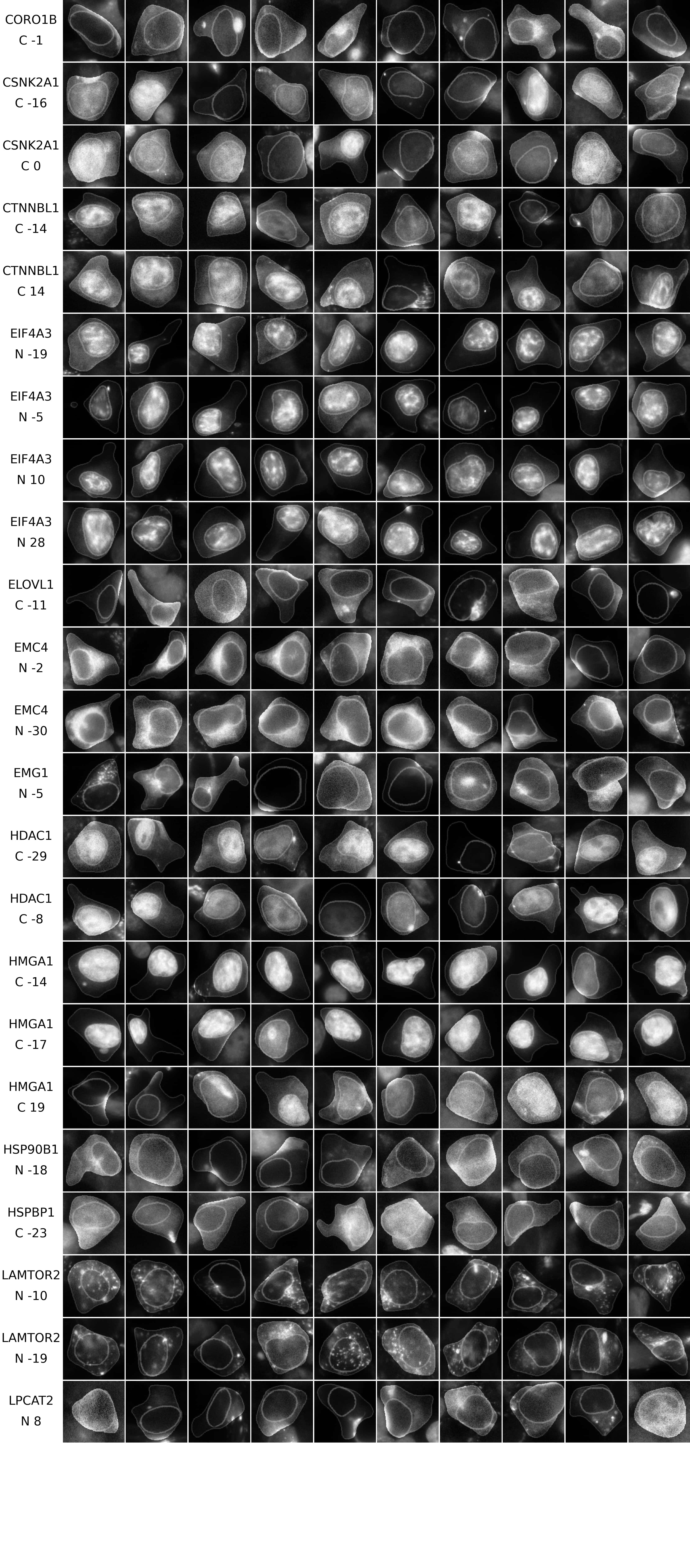

### Figure S13

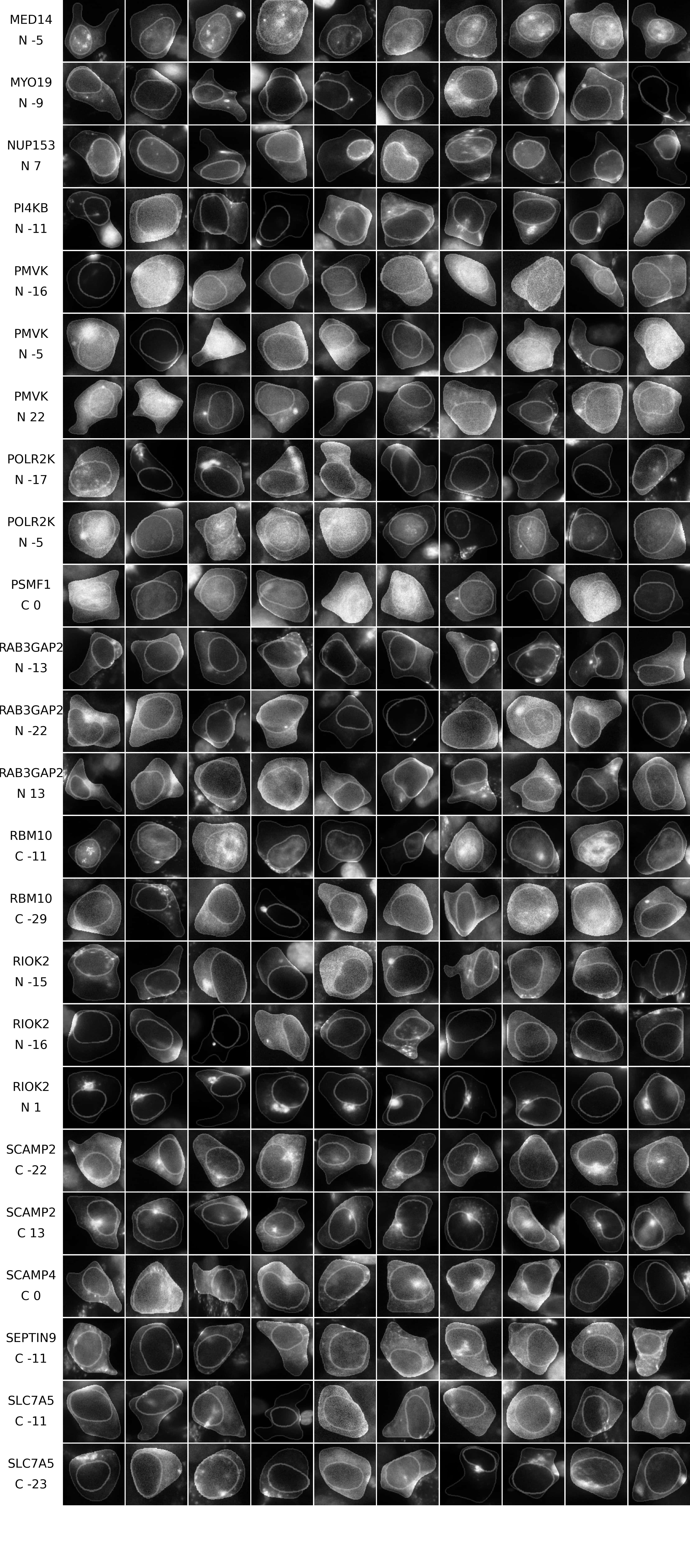

### Figure S14

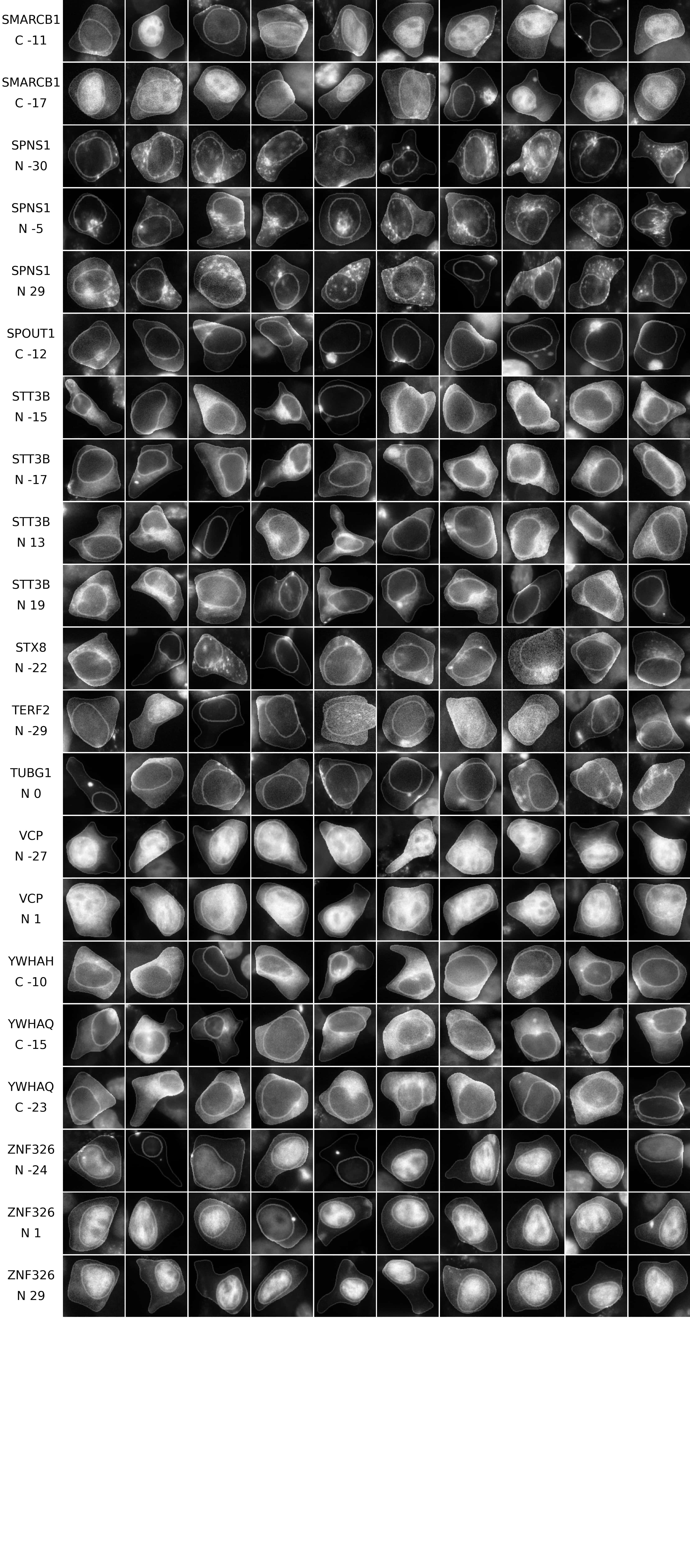

### Figure S15

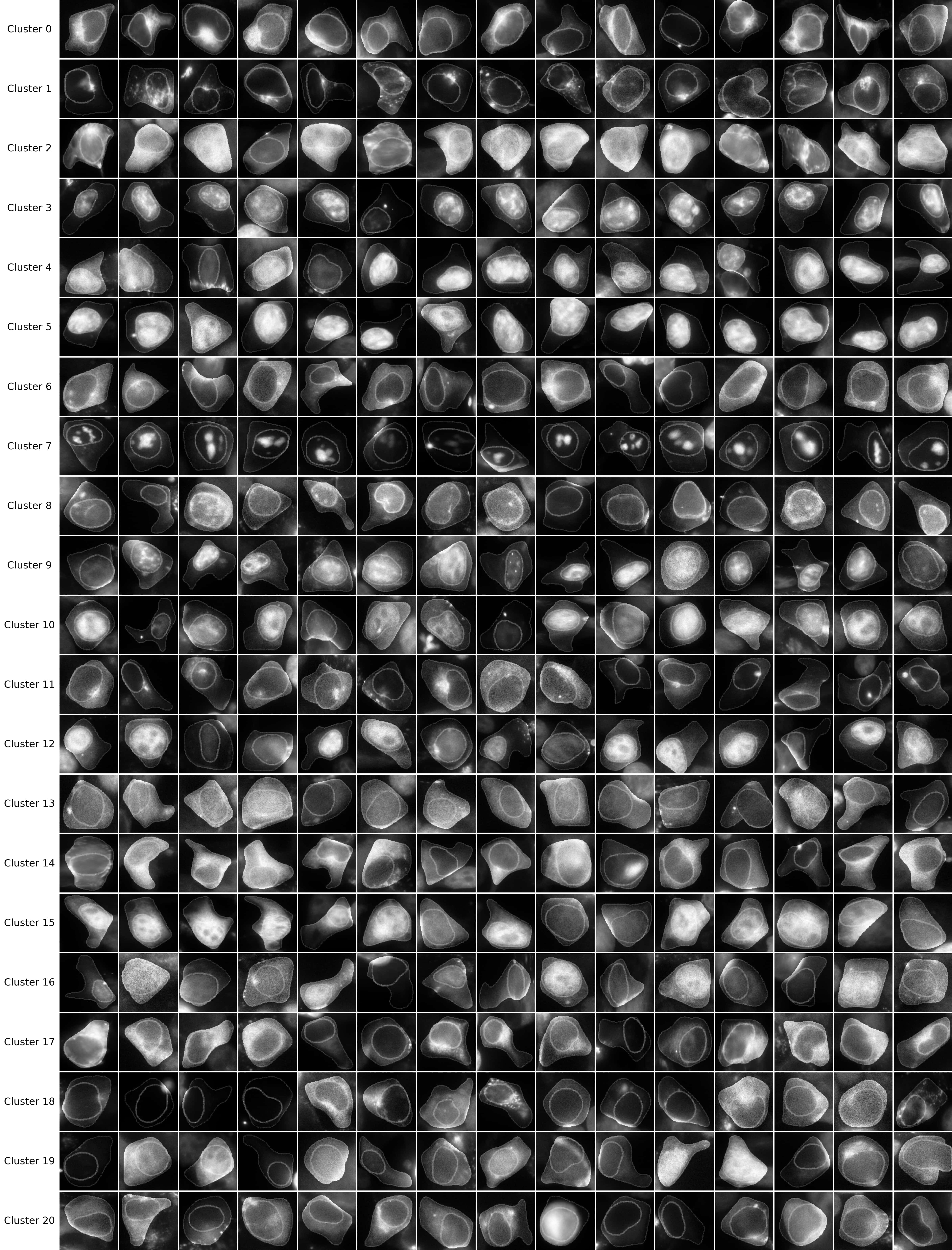
